# Sequential intra-articular HCAd-NFκB-IL-1Ra delivery improves therapeutic efficacy in post-traumatic osteoarthritis

**DOI:** 10.64898/2026.07.27.740818

**Authors:** Jiansen Yan, Zelong Dou, Camilla Fernanda Majano, Racel Cela, Ming-Ming Jiang, Syed Shayan Mehdi, Joud Al Azaat, Duncan Robert Crosby, Amanda Rosewell Shaw, RE-JOIN Consortium Investigators, Lisa Ann Yuva, Surabi Veeraragavan, Oscar Eugenio Ruiz, Donna J. Palmer, Philip Ng, Nele A. Haelterman, Masataka Suzuki, Yangjin Bae, Brendan H. L. Lee

**Affiliations:** Department of Molecular and Human Genetics, Baylor College of Medicine, Houston, TX 77030, USA; Center for Cell and Gene Therapy, Baylor College of Medicine, Texas Children’s Hospital, Houston, TX 77030, USA

**Keywords:** Osteoarthritis, high-capacity adenovirus, serotype switching, repeat injection, IL-1Ra

## Abstract

Osteoarthritis is the most common joint disease for which disease-modifying therapies remain unavailable. Intra-articular gene delivery of interleukin-1 receptor antagonist (IL-1Ra) using high-capacity adenovirus (HCAd) has shown therapeutic promise; however, the duration of therapeutic benefit and the feasibility of repeat dosing under anti-adenoviral immunity remain unresolved. Using the murine anterior cruciate ligament transection model of osteoarthritis, we show that a single intraarticular injection of HCAd5-NFκB-IL-1Ra provides structural preservation and functional improvement in early-stage osteoarthritis but fails to sustain cartilage protection as disease progresses. However, HCAd5 transduction following repeated treatment is limited due to pre-existing immunity against this serotype. Notably, exchanging serotypes for repeated treatments effectively restores vector transduction and transgene expression in both healthy and osteoarthritic joints. Leveraging this strategy, we demonstrate that sequential intra-articular HCAd-NFκB-IL-1Ra administration does not further improve pain or motor function compared to the initial treatment, but preserves cartilage as assessed by histopathology and phase-contrast μCT, irrespective of serotype. These findings establish HCAd serotype switching as a feasible approach to overcome immune barriers to repeat intra-articular gene therapy. Importantly, sequential HCAd-NFκB-IL-1Ra administration enhances therapeutic durability in post-traumatic osteoarthritis, providing a translational framework for repeat intra-articular gene delivery strategies aimed at long-term disease modification in osteoarthritis.

## Introduction

Osteoarthritis (OA) is a chronic degenerative joint disease characterized by progressive cartilage loss, synovial inflammation, and subchondral bone remodeling, and is a leading cause of chronic joint pain and disability worldwide^1^. A substantial body of evidence indicates that inflammatory signaling plays a key role in to OA pathogenesis and disease progression, with interleukin-1 (IL-1) as a central hub driving catabolic and inflammatory processes, as well as nociceptive sensitization within the joint^2,3^. Accordingly, the endogenous IL-1 receptor antagonist (IL-1Ra) has emerged as a compelling therapeutic target for OA. However, clinical efforts targeting the IL-1 pathway, typically through intra-articular delivery of IL-1Ra^4^ or systemic IL-1 inhibition using small molecules or neutralizing antibodies^5,6^, have shown limited efficacy, in part due to short intra-articular residence time or insufficient joint exposure. These limitations highlight the need for strategies to sustain local IL-1Ra activity within the osteoarthritic joint.

Gene therapy represents a promising strategy to overcome the pharmacokinetic limitations of intra-articular biologics by enabling sustained, localized transgene expression within the joint^7,8^. Among available vector platforms, high-capacity adenoviral (HCAd) vectors are particularly well suited for osteoarthritic applications, as they are devoid of nearly all viral coding sequences, support large transgene payloads, and achieve highly efficient transduction of joint-associated tissues, resulting in robust episomal transgene expression without genomic integration^9,10^. Building on these advantages, we previously developed an HCAd serotype 5-based IL-1Ra gene therapy platform regulated by a nuclear factor-κB responsive promoter (HCAd5-NFκB-IL-1Ra) to enable inflammation-driven transgene expression within the osteoarthritic joint. In preclinical studies, intra-articular administration of HCAd5-NFκB-IL-1Ra demonstrated therapeutic efficacy across multiple OA models, including mouse^11^, rat^12^, and equine models^13^, leading to long-term, regulated, local IL-1Ra expression and improvement in joint pathology and pain-related outcomes. Consistent with these findings, a phase I clinical trial (NCT04119687) of this platform demonstrated a favorable safety profile and clinically meaningful improvements in OA-associated pain and joint stiffness^14^, collectively supporting HCAd5-NFκB-IL-1Ra gene therapy as a promising disease-modifying approach for OA.

To date, most *in vivo* viral vector-mediated gene therapy approaches are administered as a single dose. This paradigm is supported by the relatively prolonged duration of transgene expression achievable with viral vectors, as well as by practical considerations related to immune responses, safety, and regulatory constraints associated with repeat administration^15^. Importantly, osteoarthritis is a chronic, progressive disease for which no FDA-approved disease-modifying treatment exists. In this context, the dynamic, time-dependent changes that occur in the whole joint as disease progresses may exceed even the stable, long-term expression capacity afforded by a single non-integrating viral vector administration. OA is characterized by continuous tissue remodeling and inflammatory activity within the joint. Following intra-articular viral gene delivery, transgene expression arises predominantly from synovial tissues, which represent the primary cellular targets of viral transduction within the joint^16^. Ongoing cellular turnover within the synovium is therefore expected to gradually reduce the population of transduced cells over prolonged periods. In addition, progressive cartilage erosion during OA may further diminish transgene expression through the loss of the relatively small number of transduced superficial chondrocytes. Together, these biological features, which are not unique to OA but are particularly relevant in the context of chronic diseases, suggest that a single vector administration may be insufficient to sustain therapeutic transgene levels across the full course of OA. Thus, it raises the question whether repeat intra-articular gene delivery could overcome these challenges and provide long-term efficacy.

A central challenge to repeat vector administration is the emergence of host immunity directed against viral capsid proteins, which is well characterized in the context of systemic gene delivery^17^. However, how pre-existing or treatment-induced anti-adenoviral immune responses influence the feasibility of repeat intra-articular gene transfer remains elusive. Early clinical experience with intra-articular HCAd-based gene therapy indicates that baseline anti-adenoviral immunity does not significantly limit HCAd5-NFκB-IL-1Ra’s therapeutic efficacy in OA^14^, highlighting an incomplete understanding of how pre-existing immune responses influence vector performance within the joint. Neutralizing antibody responses to adenoviral vectors are predominantly mediated by antibodies targeting exposed epitopes on the hexon capsid proteins^18^. Importantly, the antigenic determinants of the capsid protein vary substantially across adenoviral serotypes, reflecting marked sequence and structural divergence^19^. Leveraging these differences through serotype switching, defined as sequential administration of vectors derived from distinct adenoviral serotypes yet still encoding the same therapeutic transgene, represents a rational strategy to mitigate vector-specific immune recognition and enable repeat intra-articular gene delivery^20^. Unlike in a diverse human population with an even more diverse repertoire of anti-adenovirus antibodies, a primed, boost immunization strategy in rodents provides a model for maximal anti-adenovirus immune response. While the data may or may not predict the typical response in the human population, it serves as a potential outcome in the context of maximized pre-existing immunity.

In this study, we systematically examined the role of antiviral immunity in intra-articular adenoviral gene delivery and explore the feasibility of repeat joint administration of HCAd vectors after pre-immunization. First, we investigated how pre-existing and treatment-induced anti-adenoviral immune responses influence vector transduction following intra-articular delivery. Second, we evaluated the impact of adenoviral serotype switching on vector performance in the presence of pre-existing antiviral immunity. Finally, we examined whether repeat intra-articular administration of HCAd-NFκB-IL-1Ra can improve therapeutic efficacy in a post-traumatic osteoarthritis (PTOA) model. Our results demonstrate that in mouse, antiviral immune responses substantially restrict repeat intra-articular delivery of the same adenoviral vector, whereas serotype switching bypasses this prior immune response and enables effective vector re-administration. Importantly, repeat intra-articular administration of HCAd-NFκB-IL-1Ra enhances and prolongs therapeutic efficacy in PTOA.

## Results

### Single intra-articular administration of HCAd5-NFκB-IL-1Ra delays but does not prevent OA progression

To define the therapeutic duration following a single intra-articular HCAd5-IL-1Ra administration, mice subjected to anterior cruciate ligament transection (ACLT) received a single intra-articular injection of HCAd5-NFκB-IL-1Ra and were evaluated at 6 and 10 weeks post-surgery to assess changes in structural and functional outcomes over time. Safranin-O/Fast Green staining at 6 weeks post-ACLT revealed proteoglycan loss in both HCAd5-empty and HCAd5-IL-1Ra treated joints, while cartilage surfaces remained largely intact with minimal erosion. At this stage, HCAd5-IL-1Ra treated joints showed greater cartilage thickness compared with HCAd5-empty controls, but OARSI scores were not significantly different between groups (Fig. 1A, B). By 10 weeks post-ACLT, more advanced articular cartilage degeneration was evident in HCAd5-IL-1Ra treated joints, including focal cartilage erosions and surface discontinuities, accompanied by an increase in OARSI scores compared to 6 weeks. Synovial pathology was evaluated in parallel using H&E staining. At 6 weeks post-ACLT, synovitis scores did not differ between HCAd5-empty and HCAd5-IL-1Ra treated joints. In contrast, synovial hyperplasia was more pronounced at 10 weeks, indicating progressive synovial inflammation over time despite initial HCAd5-IL-1Ra treatment (Fig. 1A, C).

**Figure 1.**
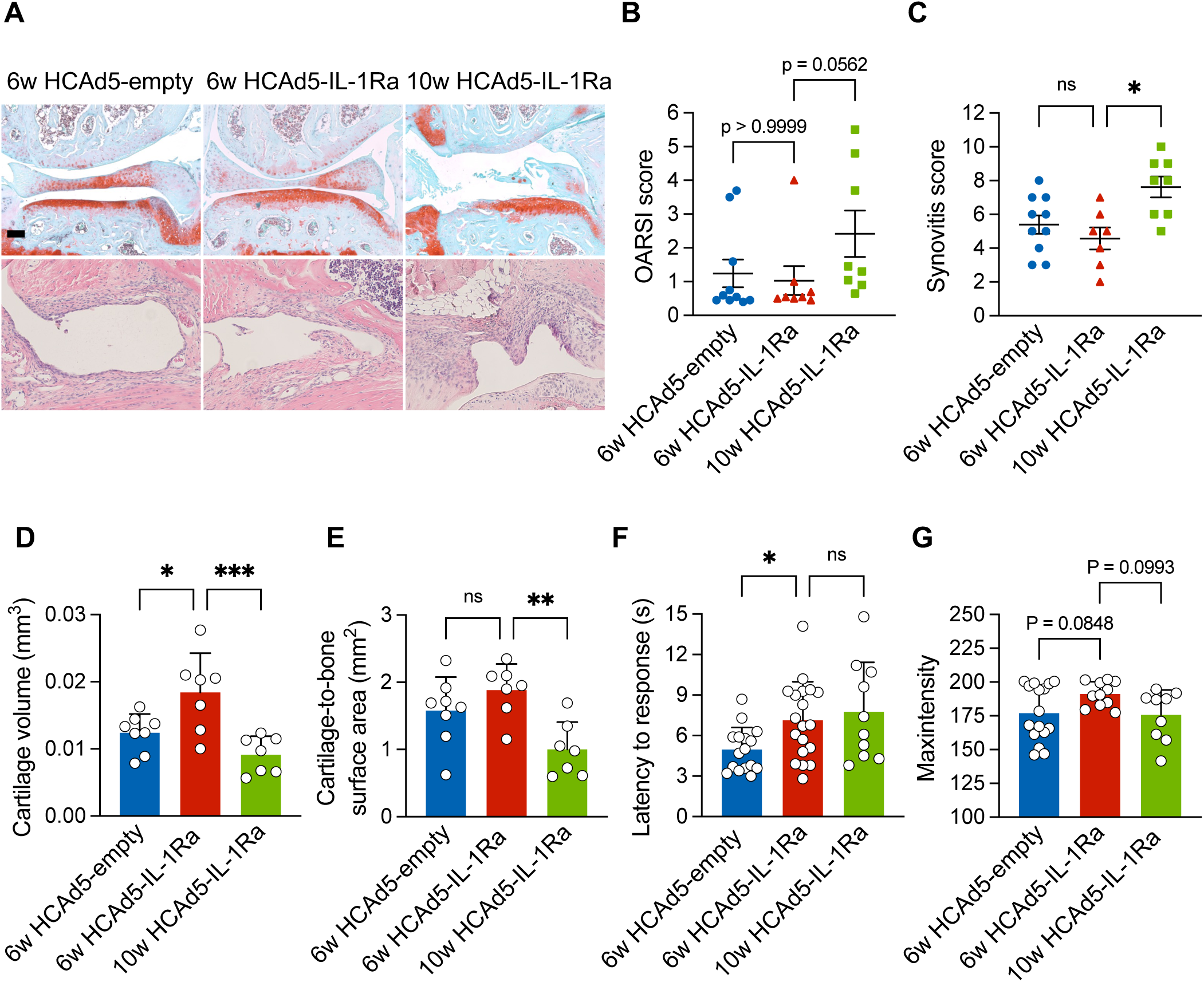
Structural, thermal nociceptive, and motor function outcomes following a single intra-articular injection of HCAd5-NFκB-IL-1Ra in an ACLT mouse model. **(A)** Representative Safranin O/Fast Green–stained sections of knee joints from mice receiving intra-articular injection of control HCAd5-empty or HCAd5-IL-1Ra one week after ACLT surgery. Joints were harvested at 6 or 10 weeks post-surgery. (**B)** Average OARSI scores of medial tibial and femoral cartilage (n = 8-10). **(C)** Synovitis score of the whole knee joint (n = 7-10). **(D)** Quantification of femoral medial cartilage volume assessed by phase contrast μCT (n = 6-8). **(E)** Quantification of femoral medial cartilage-to-bone surface area assessed by phase contrast μCT (n = 6-8). **(F)** Latency to response in the 55°C hot-plate test (n = 9-19). **(G)** Mean maximum intensity of bilateral hind paws measured by CatWalk gait analysis (n = 9-16). The scale bar is 500 μm. *p<0.05, **p<0.01, ***p<0.001; ns=no significance.

To complement histological analysis and further characterize cartilage structural changes over time, phase-contrast micro-computed tomography (µCT) was employed to assess three-dimensional cartilage morphology at high resolution. At 6 weeks post-ACLT, HCAd5-IL-1Ra treated joints exhibited higher cartilage volume compared to HCAd5-empty controls, whereas cartilage-to-bone surface area did not differ significantly between these two groups (Fig. 1D, E). This result is consistent with OARSI grading at this time point, in which cartilage surfaces remained largely intact despite decreased cartilage thickness in HCAd-empty group, indicating that early structural changes were dominated by cartilage thinning rather than surface erosion. By 10 weeks post-ACLT, cartilage volume and cartilage-to-bone surface area declined in HCAd5-IL-1Ra treated joints relative to the 6-week time point (Fig.1D, E), coinciding with the emergence of focal cartilage erosions on histological assessment. Together, these data indicate that IL-1Ra gene therapy delays, rather than inhibits, OA-induced structural changes over time.

Next, we assessed functional outcomes by pain and motor function tests. HCAd5-IL-1Ra treated mice showed significantly increased withdrawal latency in the hot plate test compared to HCAd5-empty controls at 6 weeks post-ACLT, indicating reduced thermal hyperalgesia, and this analgesic effect persisted at 10 weeks (Fig. 1F). Motor function was assessed using CatWalk gait analysis. HCAd5-IL-1Ra treatment improved motor performance, as reflected by increased maxintensity at 6 weeks post-ACLT compared to HCAd5-empty controls. However, maxintensity declined by 10 weeks relative to the 6-week time point, indicating delayed motor function decline that was not arrested by gene therapy (Fig. 1G).

Together, these data show that a single intra-articular administration of HCAd5-NFκB-IL-1Ra provides preservation of cartilage thickness and volume, accompanied by reduced pain and improved motor function. However, the structural defects and motor dysfunction are delayed rather than inhibited, suggesting that repeat intra-articular administration of HCAd5-NFκB-IL-1Ra should be investigated for improved long-term efficac

### Prior immunization with a first-generation adenovirus serotype 5 (FGAd5) impairs intra-articular HCAd5 transduction and transgene expression

Before performing repeat HCAd-NFκB-IL-1Ra administration, we characterized the impact of systemic preexisting anti-adenoviral immunity on vector transduction in joint space after intra-articular HCAd injection. To induce a robust immune response against adenovirus serotype 5, mice received two sequential intramuscular injections of a FGAd5 vector. Following this immunization, a HCAd5-CMV-β-galactosidase (LacZ) vector was administered intra-articularly (Fig. 2A). Serum neutralizing antibody assays confirmed successful induction of strong anti-adenovirus serotype 5 immunity, with a significantly higher neutralization percentage detected compared to the non-immunized group (Fig. 2B). Following intra-articular injection of HCAd5-LacZ, quantitative PCR analysis of joint derived DNA revealed a significant reduction in HCAd5 viral DNA copy number in knee joints of pre-immunized animals compared to un-immunized animals (Fig. 2C). Consistent with reduced vector genome copy numbers, lacZ mRNA levels were significantly decreased in joints from animals with preexisting immunity (Fig. 2D), accompanied by a corresponding reduction in transgene protein expression as assessed by X-gal staining of joint sections (Fig. 2E). Together, these results demonstrate that induction of systemic anti-adenoviral immunity profoundly impairs subsequent intra-articular HCAd transduction and transgene expression, in spite of local delivery to the knee joint.

**Figure 2.**
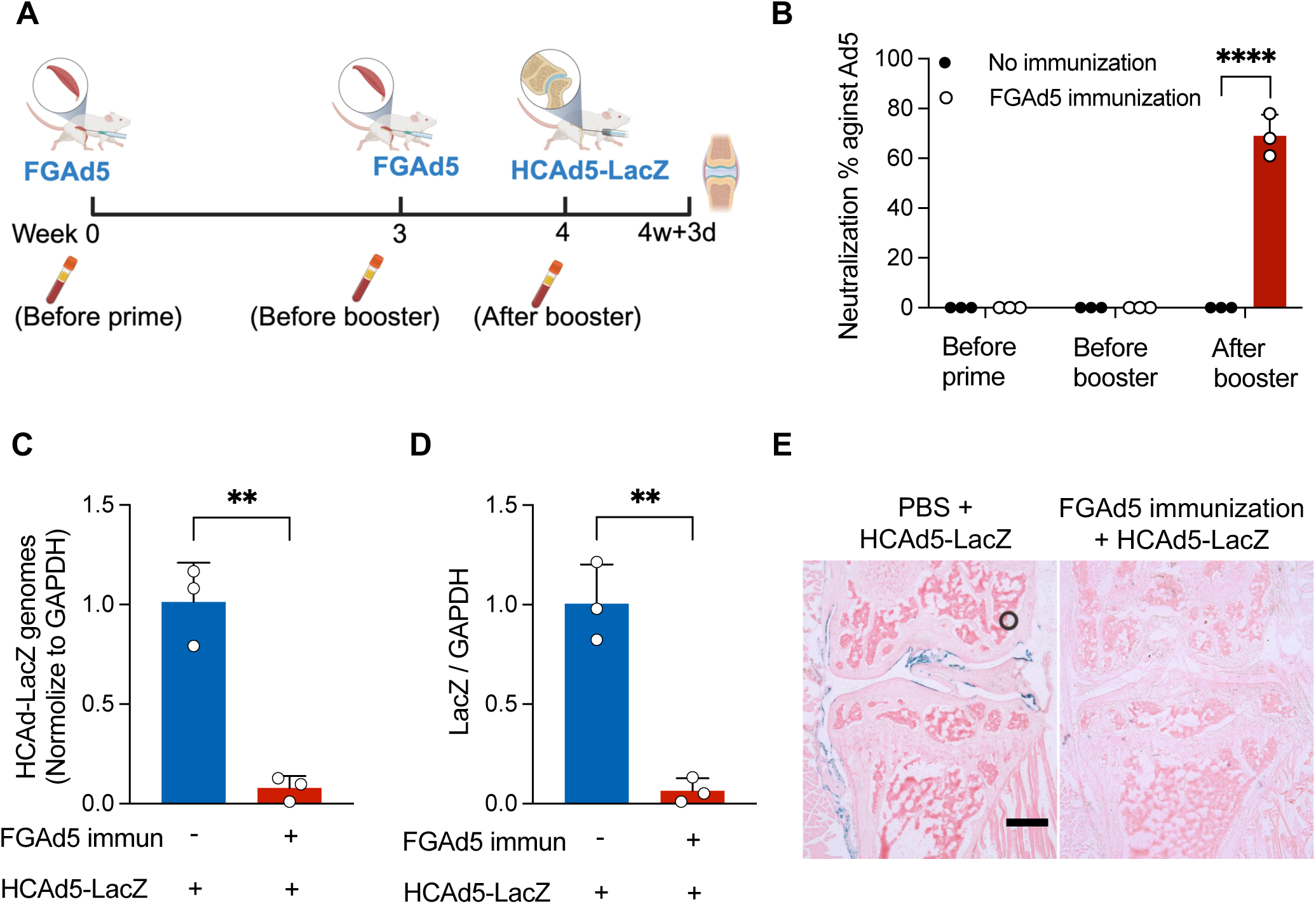
Vector genomic DNA and transgene expression following intra-articular injection of HCAd5-LacZ in FGAd5-immunized mice. **(A)** Experimental design illustrating prime and booster intramuscular injections of FGAd5 to induce pre-existing anti-Ad5 immunity, followed by intra-articular injection of HCAd5-LacZ. **(B)** Neutralization percentage of Ad5-specific neutralizing antibodies measured in serum prior to the prime injection, prior to the booster injection, and prior to intra-articular injection in control and FGAd5-immunized groups (n = 3). **(C)** Quantitative PCR analysis of viral genomic DNA from whole-joint DNA extracts (n = 3). **(D)** Quantitative PCR analysis of LacZ mRNA from whole-joint RNA extracts (n = 3). **(E)** Representative images of X-gal stained knee joints. The scale bar is 500 μm. “immun” indicates immunization. **p<0.01, ****p<0.0001.

### Alternative HCAd serotypes display comparable tissue tropism and transgene expression profiles in the knee joint compared to HCAd5

Given that induction of systemic anti-Ad5 immunity limits subsequent intra-articular HCAd5 transduction, we next evaluated whether alternative HCAd serotypes could bypass the preexisting immunity to FGAd5. Alternative HCAd serotypes were selected from the same adenoviral subgroup as HCAd5 (including HCAd1, HCAd2, and HCAd6) to minimize differences in transfection while enabling immune evasion through distinct capsid proteins. Neutralizing antibody assays revealed that FGAd5 immunization elicited serotype-specific neutralization of HCAd5, with no detectable cross-reactivity against HCAd1, HCAd2, or HCAd6 within the same subgroup (Fig. S1).

We next characterized joint tissue tropism of these alternative serotypes following intra-articular delivery in non-immunized animals. HCAd-CMV-LacZ vectors of each serotype were delivered via intra-articular injection, and transgene expression was evaluated 3 days later. X-gal staining revealed robust and localized transduction within the knee joint across all tested serotypes, with patterns comparable to HCAd5 injection. Transgene expression was observed predominantly in synovial lining and subintimal cells, soft connective tissues adjacent to the cruciate ligaments, and superficial zone cells of the meniscus (Fig. 3A), indicating that all tested HCAd serotypes retain the ability to efficiently transduce joint tissues comparable to HCAd5.

**Figure 3.**
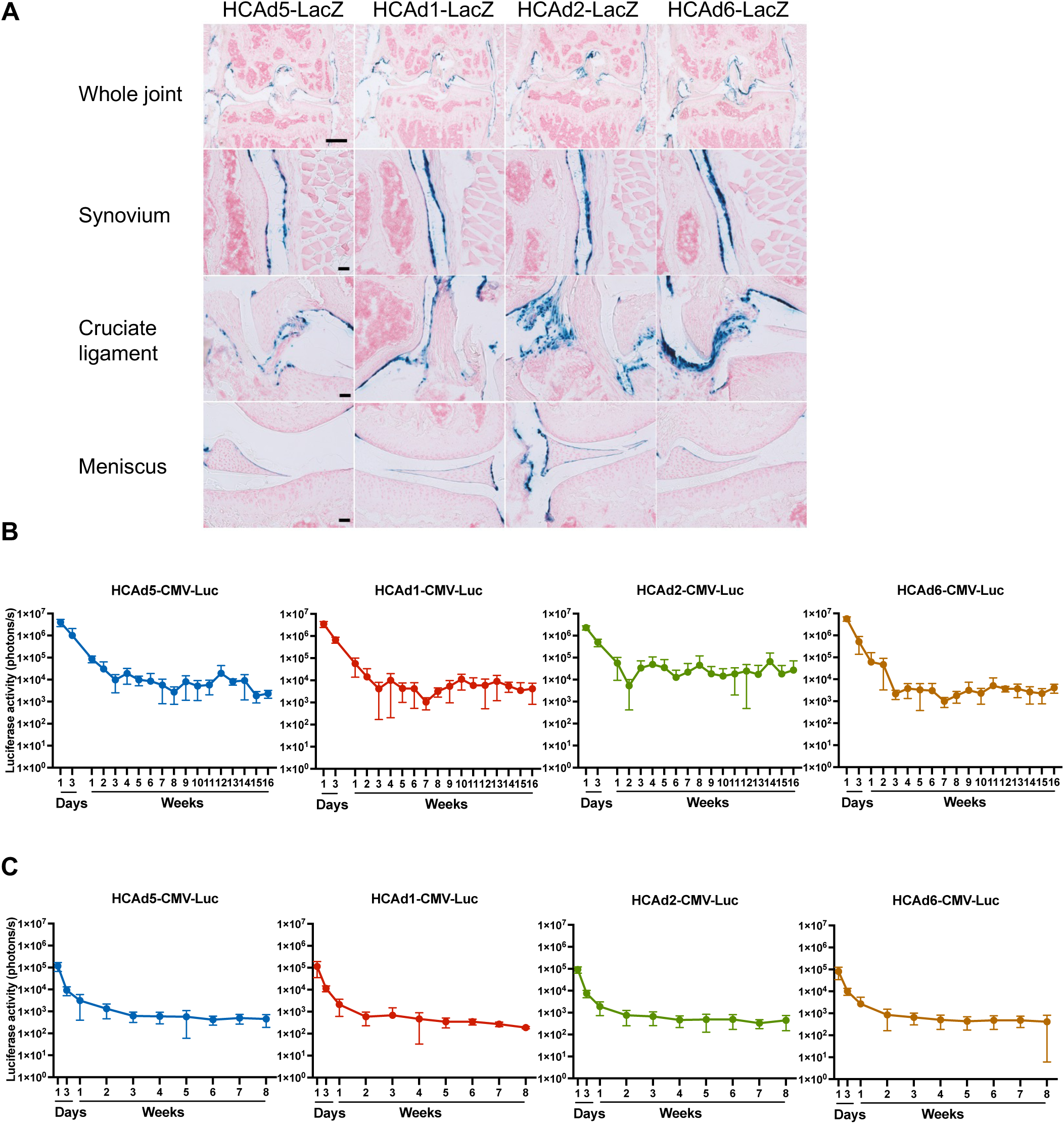
Joint tropism and transgene expression of HCAd vectors with different serotypes. **(A**) Representative X-gal staining of healthy knee joints following intra-articular injection of HCAd5-LacZ, HCAd1-LacZ, HCAd2-LacZ, or HCAd6-LacZ, demonstrating LacZ expression in the synovium, peri-ligamentous soft tissues, and the superficial zone of the meniscus. **(B)** Longitudinal bioluminescence imaging of luciferase expression in healthy knee joints injected intra-articularly with HCAd vectors of different serotypes that all express luciferase, monitored by IVIS imaging for up to 16 weeks post-injection (n = 4-8). **(C)** Osteoarthritic knee joints subjected to ACLT surgery received intra-articular injection of different HCAd serotypes expressing luciferase at 4 weeks post-surgery, and longitudinal luciferase expression was monitored by IVIS imaging for up to 8 weeks post-injection (n = 6). The scale bar for the top row is 500 μm; the scale bars for other rows are 100 μm.

To characterize transgene expression kinetics, luciferase activity was measured longitudinally following intra-articular injection of HCAd expressing luciferase from a CMV promoter (HCAd-CMV-Luc). In naïve knee joints, all serotypes exhibited a similar expression profile, with high luciferase activity detected at 24 hours post-injection, followed by a gradual decline over the subsequent 2 to 3 weeks and stabilization thereafter (Fig. 3B). Both the magnitude and temporal pattern of expression were comparable among all tested serotypes, including HCAd5. Because inflammatory status and progressive joint pathology associated with osteoarthritis could potentially influence viral transduction and transgene expression, we next evaluated these vectors’ expression kinetics in the ACLT model of OA. HCAd-CMV-Luc vectors were intra-articularly injected 4 weeks after ACLT surgery, corresponding to early-stage osteoarthritis. In osteoarthritic joints, all serotypes demonstrated expression kinetics similar to those observed in naïve joints, including an early peak at 24 hours followed by a decline and stabilization phase (Fig. 3C). No substantial differences in magnitude or duration of expression were observed between serotypes under osteoarthritic conditions, indicating that disease-associated joint structural and inflammatory changes do not alter the overall pattern of HCAd-mediated transgene expression. Together, these results demonstrate that alternative HCAd serotypes evade serotype-specific neutralizing antibodies while preserving efficient joint transduction and durable transgene expression in both healthy and osteoarthritic joints, thereby supporting serotype switching as a strategy to enable repeat intra-articular gene delivery in the face of previous immunization.

### Serotype switching restores transduction in joint tissues of FGAd5-immunized mice

We next assessed the effect of adenoviral serotype switching on intra-articular HCAd transduction under conditions of preexisting systemic immunity. Naïve mice were first immunized to induce systemic anti-adenoviral immunity and subsequently received intra-articular HCAd injection using either the same serotype (HCAd5) or an alternative serotype (HCAd1, HCAd2, or HCAd6) expressing LacZ (Fig. 4A). Neutralizing antibody assessment showed successful anti-Ad5 immunization (Fig. 4B). Vector transduction was evaluated by quantifying viral genome copy numbers within the joint. In FGAd5-immunized mice, repeat intra-articular administration of HCAd5 resulted in low viral DNA copy numbers, whereas switching to alternative serotypes markedly increased HCAd DNA levels in the joint (Fig. 4C), indicating recovery of vector transduction following serotype switching. Consistent with increased vector presence, transgene mRNA levels were elevated following serotype switching. HCAd1 and HCAd2 administration resulted in increased transgene mRNA expression compared to HCAd5 injection, with HCAd2 achieving a statistically significant increase, while HCAd6 produced a more modest elevation in mRNA levels (Fig. 4D). At the protein level, X-gal staining demonstrated robust LacZ level in joints receiving alternative serotypes, with all tested alternative serotypes showing substantial LacZ expression compared to HCAd5 (Fig. 4E). Together, these results demonstrate that adenoviral serotype switching effectively overcomes preexisting anti-adenoviral immunity and restores intra-articular HCAd transduction and transgene expression in healthy joints.

**Figure 4.**
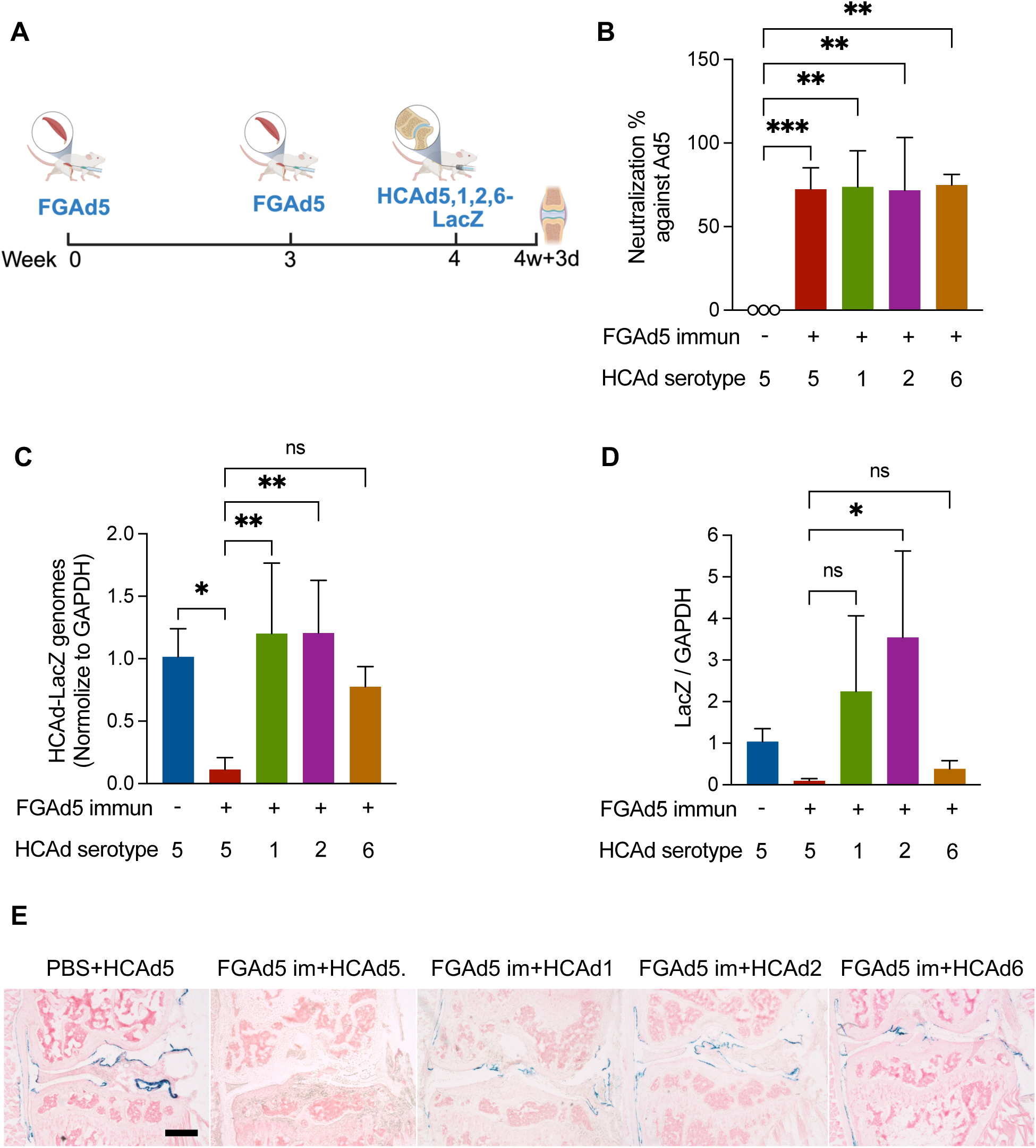
Vector transduction and transgene expression following intra-articular injection of HCAd vectors with different serotypes in FGAd5-immunized mice. **(A)** Schematic overview of the experimental design. 12-week-old mice received two intramuscular injections of FGAd5 to induce pre-existing anti-Ad5 immunity and, one week later, intra-articular injection of HCAd5-LacZ, HCAd1-LacZ, HCAd2-LacZ, or HCAd6-LacZ. Knee joints were harvested 3 days after intra-articular injection. **(B)** Neutralizing antibody assay showing the percentage of HCAd5 neutralization in serum collected after full immunization with FGAd5 (n = 3-4). **(C)** Vector genomic DNA copy numbers in whole-joint DNA extracts as assessed by quantitative PCR (n = 3). **(D)** LacZ mRNA expression measured by quantitative PCR from whole-joint RNA extracts (n = 3). **(E)** Representative images of X-gal staining of knee joints. The scale bar is 500 μm. “im” indicates immunization. *p<0.05, **p<0.01, ***p<0.001; ns=no significance.

### Serotype switching restores transduction of HCAd in osteoarthritic joints of FGAd5-immunized mice

To determine whether the effects observed in healthy joints also apply under osteoarthritic conditions, we next evaluated the effect of adenoviral serotype switching in the context of established osteoarthritis. Mice with anti-adenoviral immunity underwent ACLT surgery and subsequently received intra-articular HCAd injection using either the same serotype or alternative serotypes during early-stage OA (Fig. 5A). Serum neutralizing antibody assays confirmed robust preexisting anti-adenoviral immunity prior to ACLT surgery (Fig. 5B). In osteoarthritic joints of FGAd5-immunized mice, intra-articular administration of HCAd5 resulted in low viral DNA copy numbers, whereas switching to alternative serotypes showed HCAd DNA levels within the joint tissue that were comparable to those detected in un-immunized mouse joints (Fig. 5C). Consistent with the pattern observed in healthy joints, transgene mRNA expression was elevated following serotype switching in osteoarthritic joints. HCAd1 and HCAd2 administration increased transgene mRNA levels compared to HCAd5 injection, with HCAd2 reaching statistical significance, while HCAd6 produced a more modest increase (Fig. 5D). Together, these results demonstrate that adenoviral serotype switching restores HCAd transduction and transgene expression in osteoarthritic joints despite systemic anti-adenoviral immunity.

**Figure 5.**
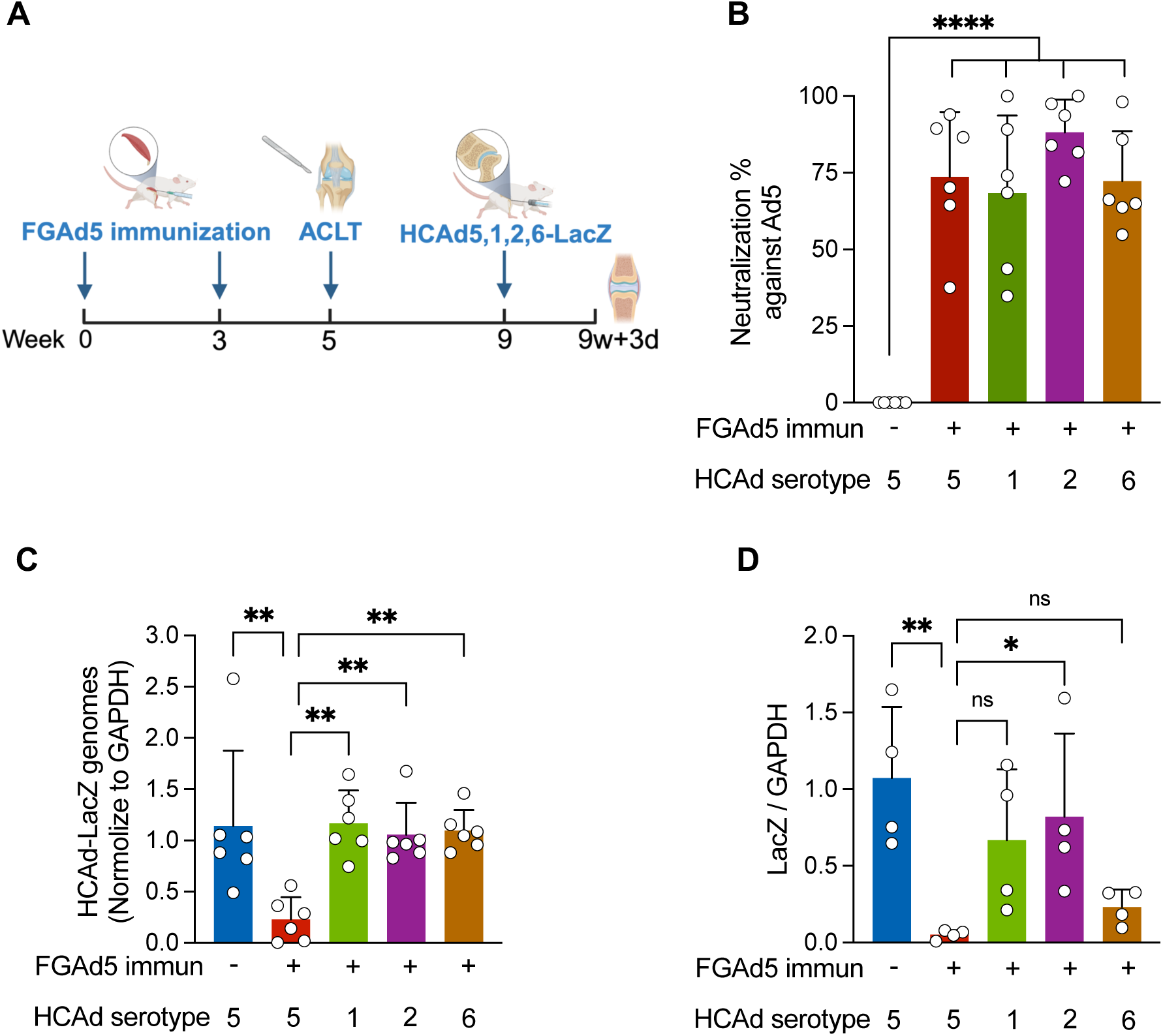
Vector transduction and transgene expression following intra-articular injection of HCAd vectors with different serotypes in FGAd5-immunized ACLT mice. **(A)** Experimental timeline. 7-week-old mice received two intramuscular immunizations with FGAd5 that were 3 weeks apart, underwent ACLT surgery two weeks following the second immunization, and were injected intra-articularly with HCAd5-LacZ, HCAd1-LacZ, HCAd2-LacZ, or HCAd6-LacZ at 4 weeks post-surgery. Knee joints were harvested 3 days after intra-articular injection. **(B)** Neutralizing antibody assay showing the percentage of HCAd5 neutralization in serum collected prior to ACLT surgery (n = 6). **(C)** Vector genomic DNA copy numbers in whole-joint DNA extracts as assessed by quantitative PCR (n = 6). **(D)** LacZ mRNA expression measured by quantitative PCR from whole-joint RNA extracts (n = 4). “immun” indicates immunization. *p<0.05, **p<0.01, ****p<0.0001; ns=no significance.

### Repeated intra-articular injection with different HCAd serotypes restores transduction in the knee joint

Following observations on the impact of systemic immunity, we next assessed joint transduction following repeated intra-articular HCAd administration using the same or different serotypes (Fig. 6A). Interestingly, a single intra-articular administration of different HCAd serotypes induced robust neutralizing antibody responses against each respective serotype. (Fig. 6B). To determine the functional consequence of this immunization on repeat dosing, HCAd transduction was quantified by measuring viral DNA copy numbers using primers specific for the lacZ transgene following a second intra-articular injection with HCAd5-LacZ, 16 weeks after the initial injection. Repeat delivery of same serotype resulted in reduced HCAd DNA copy numbers within the knee joint, indicating impaired vector transduction in the context of immunity stimulated by prior intra-articular injection (HCAd5 + HCAd5 group in Fig. 6C). In contrast, switching to a different HCAd serotype significantly increased viral DNA levels (Fig. 6C). Restoration of vector transduction by serotype switching was further supported at the protein level. X-gal staining demonstrated minimal transgene expression following repeat administration of the same serotype, whereas LacZ expression was observed in joints receiving a different HCAd serotype for the second injection (Fig. 6D).

**Figure 6.**
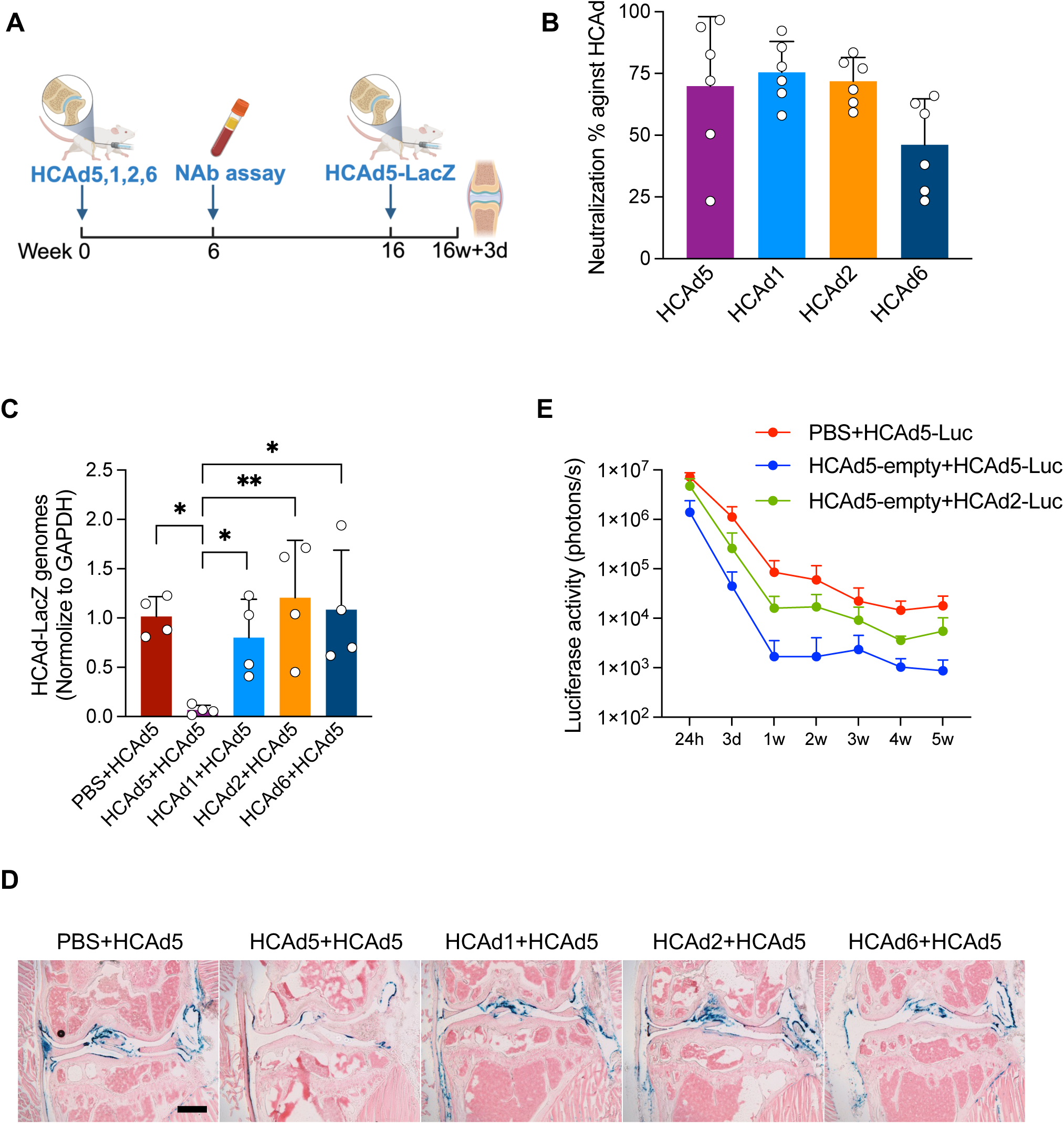
HCAd genomic DNA and LacZ expression in knee joints following repeat intra-articular injection. **(A)** Experimental timeline. 12-week-old mice received an initial intra-articular injection with HCAd5, HCAd1, HCAd2, or HCAd6, followed 16 weeks later by a second intra-articular injection with HCAd5-LacZ. Knee joints were harvested 3 days after the second injection. **(B)** Neutralizing antibody assay showing the percentage of serum neutralization against the capsid proteins of each corresponding HCAd serotype, measured 6 weeks after intra-articular injection of different HCAd serotypes (n = 6). **(C)** Quantitative PCR analysis of vector genomic DNA derived from the second HCAd5-LacZ injection in whole-joint DNA extracts using LacZ primer (n = 4). **(D)** Representative X-gal staining of knee joints following repeat intra-articular injection. **(E)** Mice received an initial intra-articular injection of PBS or empty HCAd5 vector and were re-injected 4 weeks later with HCAd5-CMV-Luc or HCAd2-CMV-Luc. Luciferase expression was monitored longitudinally by IVIS following the second injection (n = 8). The scale bar is 500 μm. “NAb” indicates neutralizing anbitody. *p<0.05, **p<0.01.

Because HCAd2 showed consistent performance across prior experiments, we next examined transgene expression dynamics following repeated intra-articular administration using a serotype switching approach. Longitudinal luciferase imaging showed that repeat intra-articular administration of the same HCAd5 serotype in healthy knee joints resulted in reduced transgene expression at all measured time points compared with the single HCAd5-Luc injection (Fig. 6E). In contrast, switching to HCAd2-Luc for the repeat injection partially restored luciferase expression over time relative to repeat administration with the same serotype, indicating that the enhancement of transgene expression conferred by serotype switching is maintained longitudinally.

### Repeated intra-articular injection with different HCAd serotypes partially restores transduction in osteoarthritic joints

We next evaluated repeat intra-articular HCAd administration in osteoarthritic joints to assess whether serotype switching preserves vector transduction during disease progression (Fig. 7A). Following ACLT surgery, an initial intra-articular injection of different serotypes of HCAd-CMV-Luc was administered 4 weeks post-surgery, corresponding to early-stage osteoarthritis, and a second intra-articular injection of HCAd5-CMV-LacZ was delivered 12 weeks post-surgery, when disease pathology was more advanced. Consistent with prior observations, a single intra-articular administration of HCAd vectors elicited significant anti-adenoviral immune responses in ACLT mice, as assessed by serum neutralizing antibody assays (Fig. 7B). When vector transduction was evaluated after repeat dosing, quantification of viral genome persistence revealed that repeat administration with the same serotype resulted in lower HCAd DNA copy numbers, whereas switching to an alternative serotype partially restored HCAd DNA levels within osteoarthritic joints (Fig. 7C). Partial recovery of vector transduction following serotype switching was further supported at the transgene expression level. Transgene mRNA analysis and X-gal staining both demonstrated increased transgene expression in joints receiving an alternative serotype compared with repeat administration of the same serotype, although expression levels remained lower than those observed after initial dosing (Fig. 7D, E).

**Figure 7.**
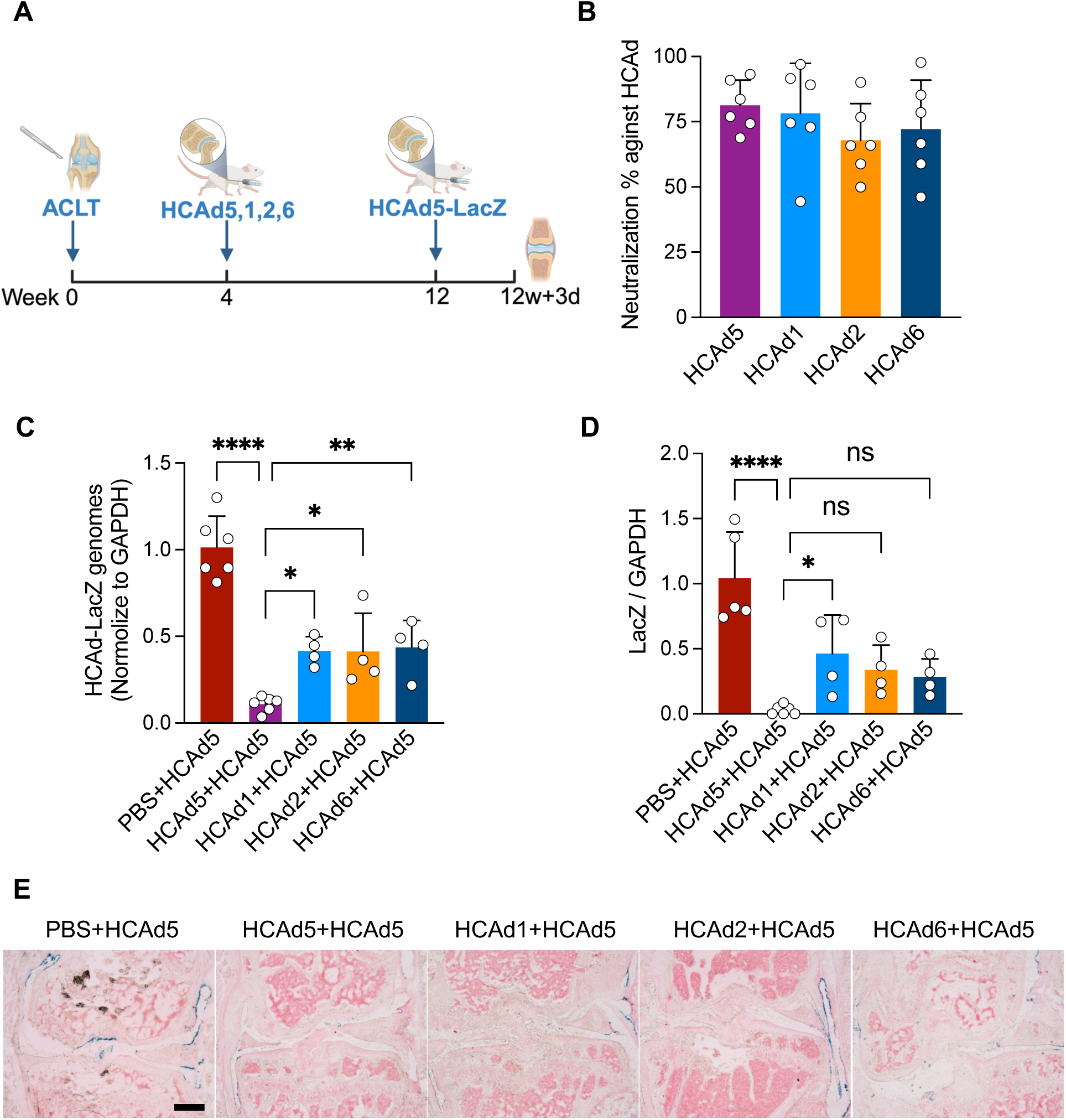
HCAd genomic DNA and LacZ expression in post-traumatic osteoarthritic joints following repeat intra-articular injection. **(A)** Experimental timeline. Mice received an initial intra-articular injection with HCAd5, HCAd1, HCAd2, or HCAd6 at 4 weeks post–ACLT surgery, which was performed at 12 weeks of age, followed by a second intra-articular injection with HCAd5-LacZ at 12 weeks post-surgery. Knee joints were harvested 3 days after the second injection. **(B)** Neutralizing antibody assay showing the percentage of neutralization against each HCAd serotype measured at 6 weeks after the first intra-articular injection (n = 6). **(C)** Quantitative PCR analysis of vector genomic DNA derived from the second HCAd5-LacZ injection in whole-joint DNA extracts (n = 4-6). **(D)** Quantitative PCR analysis of LacZ mRNA derived from the second HCAd5-LacZ injection in whole-joint RNA extracts using LacZ primer (n = 4-6). **(E)** Representative X-gal staining of knee joints following repeat intra-articular injection. The scale bar is 500 μm. *p<0.05, **p<0.01, ****p<0.0001; ns=no significance.

Together, these results demonstrate that serotype switching can partially overcome immune-mediated limitations to repeat intra-articular HCAd delivery in the context of progressive osteoarthritis, supporting the feasibility of repeat vector administration during disease progression.

### Sequential intra-articular administration of HCAd-NFκB-IL-1Ra prolongs cartilage preservation in post-traumatic OA irrespective of serotype

Following the findings on serotype switching and vector transduction, we next evaluated whether repeat intra-articular administration of HCAd-NFκB-IL-1Ra improves therapeutic efficacy in a PTOA model. We first assessed *in vitro* transgene expression levels in HEK293 cells under basal and inflammatory conditions, following transduction with two vectors of a different serotype, carrying the same transgene. ELISA revealed comparable IL-1Ra production by HCAd5-NFκB-IL-1Ra and HCAd2-NFκB-IL-1Ra at baseline. TNFα stimulation significantly enhanced IL-1Ra expression in both groups, with no difference between vectors, indicating equivalent transgene expression capacity (Fig S2).

Based on our earlier observations that HCAd5-NFκB-IL-1Ra preserves cartilage loss until 6 weeks after ACLT surgery when delivered 1 week post-surgery, a second intra-articular injection was administered at 6 weeks following the initial dose, using either the same HCAd5-NFκB-IL-1Ra or an alternative HCAd2-NFκB-IL-1Ra vector (Fig. 8A). Histological analysis confirmed that a single HCAd5-NFκB-IL-1Ra administration was not sufficient to preserve cartilage structure at 10 weeks post-injury compared to the HCAd-empty control (Fig 8B, C). In contrast, repeat intra-articular administration delayed OA progression more significantly compared to both HCAd-empty or single HCAd5-NFκB-IL-1Ra, as reflected by lower OARSI scores. Surprisingly repeated dosing maintained therapeutic efficacy regardless of whether the second dose consisted of the same serotype (HCAd5-NFκB-IL-1Ra) or a different serotype (HCAd2-NFκB-IL-1Ra) (Fig. 8B, C). Phase-contrast μCT analysis corroborated these findings, demonstrating preservation of cartilage volume and cartilage-to-bone surface area following repeat administration of either HCAd5-NFκB-IL-1Ra or HCAd2-NFκB-IL-1Ra, compared to HCAd-empty or single-dose HCAd5-NFκB-IL-1Ra (Fig. 8D, E). In the hot plate test, single-dose HCAd5-NFκB-IL-1Ra-treated mice trended towards increased withdrawal latency compared with HCAd-empty-treated animals at 10 weeks post-surgery, though this difference did not reach statistical significance (Fig. 8F). Repeat administration of either HCAd5-NFκB-IL-1Ra or HCAd2-NFκB-IL-1Ra did not further reduce thermal hyperalgesia compared to single-dose HCAd5-NFκB-IL-1Ra. Motor function was evaluated using CatWalk gait analysis. A single-dose intra-articular HCAd5-NFκB-IL-1Ra injection significantly improved maxintensity of the hind paws at 10 weeks post-injury compared with HCAd5-empty controls, whereas repeat administration of either HCAd5-NFκB-IL-1Ra or HCAd2-NFκB-IL-1Ra did not further enhance motor function beyond the effect achieved with the initial treatment regardless of serotypes at 10-week timepoint (Fig. 8G).

**Figure 8.**
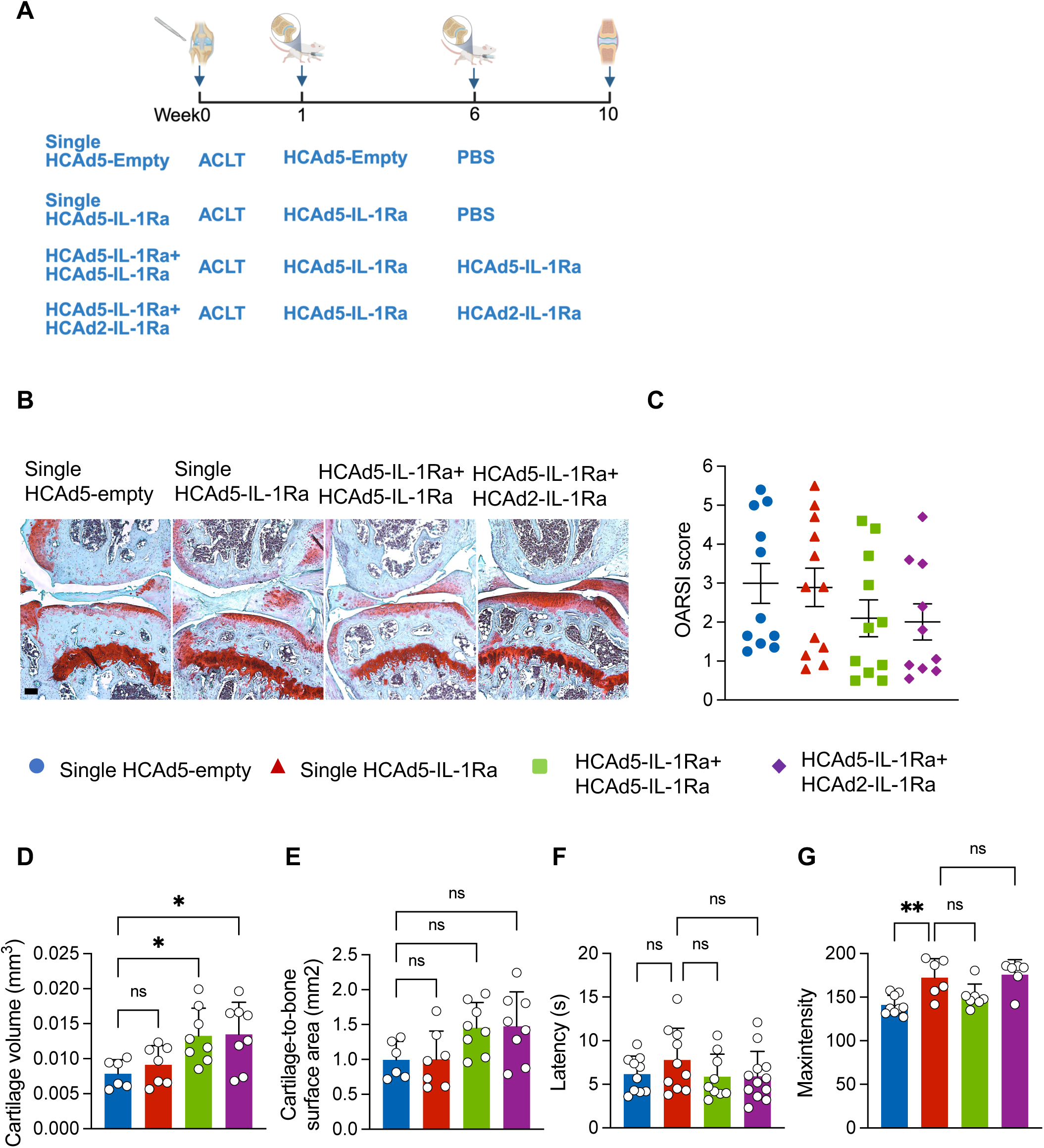
Structural, thermal nociceptive, and motor function outcomes following single or repeat intra-articular injection of HCAd-NFκB-IL-1Ra in a post-traumatic osteoarthritis model. **(A)** Schematic of the experimental design. Mice received a single intra-articular injection of HCAd5-empty or HCAd5-NFκB-IL-1Ra one week post-surgery. For repeat-injection groups, mice received an initial intra-articular injection of HCAd5-NFκB-IL-1Ra, followed by a second intra-articular injection 5 weeks later with either HCAd5-NFκB-IL-1Ra or HCAd2-NFκB-IL-1Ra. Knee joints were harvested at 10 weeks post-surgery. **(B)** Representative Safranin O/Fast Green stained sections of knee joints. **(C)** Average OARSI scores of medial tibial and femoral cartilage (n = 10-12). **(D)** Quantification of medial cartilage volume by phase contrast μCT (n = 6-8). **(E)** Quantification of medial cartilage-to-bone surface area by phase contrast μCT (n = 6-8). **(F)** Latency to response in the 55°C hot-plate test (n = 9-11). **(G)** Mean maximum intensity of bilateral hind paws measured by CatWalk gait analysis (n = 6-9). The scale bar is 100 μm. *p<0.05, **p<0.01; ns=no significance.

Together, these results demonstrate that sequential intra-articular administration of HCAd-NFκB-IL-1Ra prolonged structural protection in PTOA independent of serotype choice, while no further improvement was observed in pain or motor function following repeat administration.

## Discussion

*In vivo* viral gene therapy has long been constrained by concerns regarding the safety and feasibility of repeat vector administration, particularly for platforms in which anti-capsid immunity limits redosing^15,17^. In this study, we demonstrate that repeat intra-articular delivery of HCAd vectors is feasible in the knee joint when immune barriers associated with anti-adenoviral immunity blocks re-transduction. We show that anti-adenoviral immune responses, whether arising from classical prime-boost intramuscular administration or elicited by prior intra-articular delivery, substantially impair subsequent HCAd transduction in the joint. Importantly, we identify adenoviral serotype switching as an effective strategy to restore vector transduction and transgene expression in both healthy and osteoarthritic joints. Leveraging this approach, we further demonstrate that while the therapeutic effects of a single intra-articular HCAd5-NFκB-IL-1Ra administration diminish over time in a PTOA mouse model, sequential intra-articular dosing extends protective effects during disease progression. Together, these findings establish repeat intra-articular HCAd administration as a tractable challenge rather than a fundamental limitation and highlight serotype switching as a practical strategy for extending the effects of therapeutic gene therapy in osteoarthritis.

A key finding of this study is that pre-existing anti-adenoviral immunity elicited by immunization can be a barrier to repeat intra-articular HCAd administration. Immune constraints on vector redosing have been widely documented across viral gene therapy platforms, notably for adeno-associated virus and first generation adenovirus, where neutralizing antibodies and capsid-specific immune responses restrict repeat dosing and have driven the development of strategies such as serotype switching^20,21^, capsid engineering^22,23^, and immunomodulatory strategies^24^. Consistent with these broader principles, our data demonstrate that both systemic anti-adenoviral immunity and localized immune responses elicited by intra-articular HCAd injection are sufficient to reduce subsequent joint transduction of HCAd in mice. Notably, our data show that even a single intra-articular HCAd administration robustly induces systemic neutralizing antibody generation that significantly limits repeated dosing using the same serotype, indicating that the joint is not as immunologically silent of an environment as it was previously considered^25^. By directly examining the effects of preexisting immunity and serotype switching in healthy knee joints, and further extending this analysis to osteoarthritic joints, our study defines the magnitude and functional consequences of anti-capsid immunity in a disease-relevant intra-articular setting. These findings establish serotype switching as an effective strategy for enabling repeat intra-articular HCAd gene delivery and demonstrate that immune constraints on vector redosing also apply to HCAd gene therapy in the knee joint.

The therapeutic effects observed with repeat intra-articular HCAd-NFκB-IL-1Ra administration provide important insights into both osteoarthritis biology and the factors that contribute to durable gene therapy efficacy. Our data demonstrate that a single intra-articular delivery of HCAd-NFκB-IL-1Ra delays cartilage degeneration early after injury, but that effect diminishes as disease progresses. In contrast, repeat intra-articular administration extends the vector’s therapeutic window, further delaying cartilage loss during a critical window of post-traumatic osteoarthritis progression, indicating that reinforcement of IL-1Ra delivery can extend the structural benefits of gene therapy^26^. Surprisingly, prolonged cartilage preservation was observed following repeat dosing regardless of whether the second injection used the same or a different adenoviral serotype. This finding indicates that therapeutic benefit may not require complete restoration of vector transduction to levels observed after the initial administration at least for this vector and transgene. Rather, partial or heterogeneous transgene expression following repeat dosing appears sufficient to achieve biologically meaningful effects on joint structure at least in this preclinical surgical model.

Osteoarthritis progression reflects the cumulative impact of inflammatory and catabolic processes over time^27^, and repeated reinforcement of IL-1Ra expression may therefore shift the local joint environment toward a less destructive state, even when overall vector persistence is reduced. Consistent with this concept, emerging clinical data from a phase I study of HCAd-NFκB-IL-1Ra (PCRX-201) demonstrate sustained improvements in pain and function for up to three years following a single intraarticular injection^14^. Notably, pre-existing anti-Ad5 neutralizing antibodies did not impact pain or stiffness outcomes in patients, suggesting that complete transduction may not be required to achieve meaningful therapeutic benefit.

An additional factor likely contributing to this phenomenon is the use of an inflammation-responsive NF-κB promoter to drive IL-1Ra expression. Under this regulatory scheme, transgene output is influenced not only by vector genome copy number but also by the inflammatory state of the joint microenvironment^13,28^. As joint pathology progresses rapidly between 6 and 10 weeks after injury, increased inflammatory signaling may enhance NF-κB dependent transgene expression in successfully transduced cells following repeat administration. This disease state-coupled regulatory feature may provide a plausible explanation for why repeat intra-articular delivery, including same-serotype redosing, can sustain structural protection despite reduced vector transduction efficiency. In contrast to structural outcomes, repeat HCAd-NFκB-IL-1Ra administration did not further improve pain-related behaviors beyond those achieved with a single injection. Notably, a single HCAd5-NFκB-IL-1Ra administration was sufficient to confer sustained analgesic and functional benefits over the course of disease progression, suggesting that the level and duration of IL-1Ra expression achieved after the initial dose are adequate to modulate chronic pain-related pathways. This dissociation between preservation of cartilage and pain relief is consistent with the multifactorial nature of osteoarthritis pain, in which peripheral and central pain mechanisms may be partially uncoupled from joint pathology^29^. Together, these findings suggest that repeat intra-articular HCAd-NFκB-IL-1Ra administration effectively prolongs cartilage preservation during PTOA progression, while highlighting the complexity of targeting pain and structure within a single therapeutic framework.

The ability to achieve therapeutic benefit through repeat intra-articular HCAd administration has important translational implications for both osteoarthritis therapy and adenoviral vector development. Given that osteoarthritis is a chronic and progressive disease, sustained modulation of the joint microenvironment is likely required to achieve durable benefits. Our findings demonstrate that repeat local delivery of HCAd vectors is feasible and therapeutically meaningful, even in the presence of experimentally induced pre-existing anti-adenoviral immunity, thereby addressing a central challenge that has historically limited the clinical application of adenoviral gene therapy. From a translational perspective, the observation that structural protection can be sustained following repeat dosing with either the same or a different serotype provides important flexibility for clinical development. Same-serotype redosing is particularly attractive, as it avoids the need for multiple vector platforms, simplifies manufacturing and regulatory pathways, and may reduce development costs and barriers to patient access^30^. Also, the heterogeneous nature of pre-existing immunity in the human population may allow for re-transduction in a proportion of treated patients in contrast to the experimentally induced maximal immunity observed in the murine isogenic background used for this study. At the same time, serotype switching offers an effective strategy to enhance vector transduction when immune barriers are more pronounced, effectively broadening the therapeutic window for repeat administration in the subset of human subjects for whom pre-existing immunity may be limiting.

Beyond IL-1Ra-based gene therapy, our findings have broader implications for intra-articular gene delivery strategies. Serotype switching enables sequential delivery of distinct therapeutic payloads, such as combining anti-inflammatory, anti-catabolic, or anabolic factors, which may be particularly relevant given the multifactorial nature of osteoarthritis^11,31^. More broadly, the demonstration that HCAd vectors can be repeatedly administered to the joint while maintaining therapeutic efficacy bolsters their potential as a versatile platform for chronic diseases that require sustained or adjustable gene expression over time. Finally, the flexibility of the HCAd production system allows for a simple alternative use of the helper virus serotype of choice used in production to achieve a serotype switch in the final product and can hence maintain a low cost of production which is particularly important for treating a common disease.

Several limitations of this study should be acknowledged. First, all experiments were performed in a murine PTOA model, and immune responses to adenoviral vectors differ between mice and humans. Future studies will be needed to define how preexisting immunity and repeat intra-articular dosing translate to human joints. Second, this study focused on IL-1Ra as a single therapeutic payload. Although repeat HCAd-NFκB-IL-1Ra administration prolonged cartilage preservation, pain-related outcomes were not changed despite redosing, underscoring the multifactorial nature of osteoarthritis pain. Future studies will be required to evaluate the utility of repeat dosing and serotype switching in the context of other single target or multi-target sequential gene therapy strategies^32^. Finally, the timing and frequency of repeat intra-articular administration were not systematically studied. Further work exploring disease stage-dependent dosing strategies and alternative transgene designs may enhance the durability and scope of therapeutic benefit.

In summary, this study demonstrates that repeat intra-articular administration of high-capacity adenoviral vectors is feasible and therapeutically meaningful in a post-traumatic osteoarthritis model in mice. We show that anti-adenoviral immunity, whether induced by prior intramuscular immunization or induced by prior therapeutic joint delivery, restricts subsequent vector transduction, but this barrier can be effectively mitigated through serotype switching. Importantly, sequential intra-articular delivery of HCAd-NFκB-IL-1Ra prolongs cartilage preservation during disease progression, even when complete restoration of transduction is not achieved, highlighting the importance of repeated local reinforcement of anti-inflammatory signaling in this regulated expression construct. Together, these findings establish repeat intra-articular HCAd administration as a viable strategy for chronic joint disease and provide a framework for immune-aware, adaptable gene therapy in osteoarthritis. By defining the interplay between vector immunity, serotype selection, and therapeutic durability within the joint, this work advances both the translational development of IL-1Ra based therapy and the broader application of HCAd platforms for other diseases requiring repeat gene delivery.

## Materials and methods

### Animals

All animal experiments were approved by the Institutional Animal Care and Use Committee (IACUC) of Baylor College of Medicine. Female FVB/N (RRID IMSR_JAX: #001800) mice were used for all *in vivo* studies and were purchased from The Jackson Laboratory. Mice were maintained under specific pathogen-free conditions with a 12-hour light/dark cycle and had ad libitum access to food and water. All mice were 12 weeks of age at the time of the initial experimental intervention. In studies requiring pre-immunization followed by ACLT surgery, mice received the initial FGAd5 immunizing injection at 7 weeks of age to ensure a uniform age at the time of ACLT surgery and subsequent analyses. Animals were randomly assigned to experimental groups. All behavioral, histological, and molecular outcome assessments were conducted with investigators blinded to treatment allocation.

### High-capacity adenovirus and First-generation adenovirus

HCAd vectors lacking all viral coding sequences were used for intra-articular gene delivery. Therapeutic vectors encoding IL-1Ra were generated under the control of an NFκB-responsive promoter, including HCAd5-NFκB-IL-1Ra and HCAd2-NFκB-IL-1Ra. An HCAd-empty vector lacking a transgene cassette and containing stuffer DNA was used as a control where indicated. For transduction efficiency and longitudinal transgene expression studies, HCAd reporter vectors expressing LacZ or luciferase under the control of the CMV promoter were generated in multiple serotypes, including HCAd1, HCAd2, HCAd5, and HCAd6. Within each transgene-matched vector set, HCAd vectors were identical in genomic sequence, differing only in capsid serotype. First-generation adenoviral vectors were used exclusively for systemic immunization. These vectors were E1/E3-deleted and did not encode therapeutic transgenes. All HCAd vectors were produced using a helper virus-dependent system^33^. Vector titers were determined based on viral particle concentration calculated from optical density measurements, and equivalent doses were used across serotypes within individual experiments.

### ACLT model

Post-traumatic osteoarthritis was induced using a bilateral anterior cruciate ligament transection model^34^. All surgical procedures were performed under isoflurane anesthesia adhering to aseptic techniques. Sustained analgesia was provided with subcutaneous buprenorphine (1mg/kg), and local analgesia was achieved by intracutaneous injection of lidocaine at the surgical site prior to incision. A medial parapatellar approach was used to access the knee joint. The patella was gently dislocated laterally to expose the joint cavity, and the anterior cruciate ligament was identified under a surgical microscope. The ligament was completely transected using a fine microsurgical knife. Hemostasis was carefully achieved by gentle compression of the joint tissues with an eye spear. Successful transection was confirmed by visualization of the completely severed ACL stump attached to the tibial plateau. The joint capsule was closed, and the skin was sealed using surgical tissue adhesive. Postoperatively, mice were allowed unrestricted activity and were monitored routinely for recovery and signs of distress. To further reduce postoperative pain, an additional subcutaneous dose of buprenorphine (1mg/kg) was administered 72 hours after surgery.

### FGAd5 immunization

To induce systemic anti-adenoviral immunity, mice were immunized using a first-generation adenovirus as previously described^35^. Immunization was performed by intramuscular injection into the thigh muscle at a dose of 1 × 10⁹ viral particles diluted in 100 µL sterile phosphate-buffered saline (PBS). Two immunizations were administered per mouse with a 3-week interval between injections. The first injection was delivered to the left thigh muscle, followed by a second injection into the right thigh muscle.

### Intraarticular injection of HCAd

HCAd vectors were administered by intra-articular injection into the knee joints. All injections were performed bilaterally. For each joint, 1 × 10⁹ viral particles were diluted in 5 µL sterile PBS and injected into the joint space using a microsyringe with 33G needle (Hamilton, cat# 1702). For experiments involving HCAd-NFκB-IL-1Ra, the initial intra-articular injection was performed 1 week after ACLT surgery. In studies requiring repeat dosing of HCAd-NFκB-IL-1Ra, a second intra-articular injection was administered at 6 weeks post-surgery. Injections were carried out under aseptic conditions, and mice were allowed to recover immediately after the procedure under standard housing conditions.

### Histology

Knee joints were harvested and fixed in 4% paraformaldehyde (PFA, MilliporeSigma, cat# P6148) for 48 hours at 4 °C, followed by decalcification in 10% EDTA (VWR, cat# 0322) for 10 days at 4 °C. Decalcified tissues were processed for paraffin embedding, and formalin-fixed paraffin-embedded blocks were sectioned coronally at a thickness of 7 µm. Safranin O/Fast Green staining was performed on every other two sections throughout the joint from the anterior to posterior region. Five consecutive Safranin O/Fast Green-stained sections spanning the middle to posterior joint region, where cartilage damage is most pronounced in this model, were selected for quantitative analysis. Cartilage pathology was assessed using the Osteoarthritis Research Society International (OARSI) scoring system^36^. Given that pathological changes in the ACLT model predominantly occur in the medial compartment of the knee joint, OARSI scoring was performed on the medial femoral and medial tibial cartilage. For each knee, OARSI scores were determined for both compartments on each of the five sections, averaged within each compartment, and subsequently averaged between the medial femoral and tibial cartilage to generate a single medial joint cartilage score. Synovitis was evaluated on H&E-stained sections from comparable posterior regions. Synovial hyperplasia was scored using a standardized system across all four quadrants^37^, and scores were summed to generate a total score. All section selection and scoring were performed by an investigator masked to experimental group allocation.

### Phase-contrast μCT

Phase-contrast µCT was performed as previously described to enable high-resolution, three-dimensional visualization and quantitative analysis of articular cartilage in murine knee joints^38^. Knee joints were fixed with glutaraldehyde (Thomas Scientific, cat# C762Q38) and subsequently treated with an osmium-based contrast solution to enhance cartilage contrast prior to imaging. Imaging was performed using a Bruker SkyScan 1272 X-ray μCT scanner. Three-dimensional image reconstruction and quantitative analyses were conducted using Dragonfly software. Cartilage volume and cartilage surface area were quantified based on 100 most central sagittal slices of the medial femoral condyle. All image reconstruction, section selection, and quantitative analyses were performed by an investigator masked to experimental group allocation.

### Serum neutralizing antibody assay

Neutralizing antibodies against adenovirus were quantified using a cell-based luciferase reporter assay. Blood samples were collected using standard retro-orbital bleeding techniques, and serum was isolated for downstream analyses. For neutralization assays, serum samples were diluted 1:10 in PBS and incubated with serotype-matched HCAd-CMV-Luc reporter vectors for 1 hour at 37 °C to allow antibody-virus complex formation. In selected experiments evaluating cross-serotype neutralization following FGAd5 immunization, immune sera were incubated with HCAd reporter vectors of different serotypes under the same conditions. PBS mixed with virus in the absence of serum was used as a negative control to define maximal transduction. Serum-virus mixtures were then added to A549 cells seeded in 96-well plates and incubated at 37 °C for 30 minutes, followed by replacement with fresh culture medium. Plates were incubated overnight at 37 °C. Cells were lysed the following day using Reporter Lysis Buffer (Promega, cat# E4030), and plates were subjected to a freeze-thaw cycle at −80 °C to ensure complete cell lysis. Luciferase activity was then measured using the Luciferase Assay Reagent (Promega, cat# E1483) according to the manufacturer’s instructions. Luciferase signals were quantified using a standard curve generated by infecting cells with defined amounts of HCAd-CMV-Luciferase reporter virus in the absence of serum. Linear regression analysis was performed to calculate relative luciferase activity. Neutralization activity was determined by normalizing luciferase signals to PBS-treated virus-only controls.

### Hot plate test

Mice were acclimated to the testing room for 30 min prior to assessment. All experiments were performed in the morning, with testing conducted at the same time of day across different experimental batches to minimize circadian variability. Testing was conducted under standard room lighting conditions without white noise or additional auditory stimulation. Mice were placed individually on a heated metal surface (Hot Plate Analgesia Meter; Columbus Instruments) maintained at 55 °C, and behavior was recorded by video. The latency to a nociceptive response was determined during offline analysis. Nociceptive responses were defined as hind paw shaking, hind paw licking, or jumping, regardless of laterality. A cutoff time of 15 seconds was applied to prevent tissue injury. Mice that did not exhibit a response within this period were removed from the plate and assigned the cutoff latency. All behavioral assessments and analyses were performed by an investigator masked to experimental group allocation.

### CatWalk test

Gait and motor function were assessed using an automated CatWalk gait analysis system (CatWalk XT, version 10.6; Noldus Information Technology). Testing conditions were consistent with those used for hot plate test, except that room lighting was turned off during gait assessment. Mice were allowed to voluntarily traverse the glass walkway, and paw placement and gait parameters were recorded using the integrated high-speed camera system. Five compliant runs were collected for each mouse, defined as uninterrupted crossings with consistent locomotion. Runs that did not meet these criteria were excluded from analysis. Gait parameters were analyzed using the manufacturer’s software. Maximum intensity was used as the primary outcome measure and was defined as the peak paw contact intensity during stance. For each mouse, maximum intensity values were first automatically averaged by the CatWalk software across the five compliant runs for each hind limb. The resulting left and right hind limb values were then averaged by the analyzer to generate a single maximum intensity value per mouse for statistical analysis. All data acquisition was performed by an investigator masked to experimental group allocation.

### DNA, RNA extraction and purification and qPCR

Knee joints were harvested and carefully dissected to remove all surrounding muscle tissue. Joint-associated soft tissues, including the joint capsule, synovium, menisci, and cruciate ligaments, were retained for analysis. Articular cartilage and subchondral bone were excluded, as they are not major target tissues for HCAd transduction^16^. Dissected tissues were snap-frozen in liquid nitrogen immediately after collection and stored at −80 °C until processing. Tissues were homogenized in 1 mL TRIzol reagent (Thermo Fisher Scientific, cat# 15596026) per sample using a tissue homogenizer (Qiagen) with two cycles at 30 Hz for 1 minute each. Following phase separation, DNA and RNA were simultaneously extracted from the appropriate fractions using standard TRIzol-based protocols. RNA was further purified using a column-based cleanup kit (Zymo Research, cat# R2062) to improve purity and remove residual contaminants. Purified RNA was reverse transcribed into cDNA using the iScript cDNA synthesis kit (Bio-Rad, cat# 1708890). qPCR was performed using SYBR Green (Roche, cat# 04887352001) based real-time PCR to quantify gene expression and vector genome levels. Quantitative PCR was performed using gene-specific primers. For analysis of transgene expression, RT-qPCR was performed using LacZ primers, with GAPDH used as the reference gene for normalization. For vector genome copy number analysis, quantitative PCR was performed on genomic DNA using the same LacZ primers, and values were normalized to genomic GAPDH. The LacZ primer sequences were GTATCGCCAAAATCACCGCC and CGTTTCGTCAGTATCCCCGT. GAPDH primers were CGACCCCTTCATTGACCTCAACT and GGCCTCACCCCATTTGATGTTAG. Genomic GAPDH primers were ACCACAGTCCATGCCATCAC and TCCACCACCCTGTTGCTGTA. All reactions were run in technical replicates, and data were analyzed using standard comparative threshold cycle methods.

### ELISA

HEK293 cells were seeded in 6-well plates at 1.5 × 10^5^ cells per well 24 h prior to transduction. Cells were transduced with HCAd5-NFκB-IL-1Ra or HCAd2-NFκB-IL-1Ra at an MOI of 100 viral particles per cell. At 24 h post-transduction, recombinant human TNFα (25 ng/mL, R&D Systems, cat# 210-TA) was added, and supernatants were collected 48 h later for IL-1Ra quantification by an ELISA kit (R&D System, cat# MRA00) according to the manufacturer’s instructions.

### Luciferase assay

Bioluminescence imaging was performed using an IVIS Lumina II system (Revvity) for experiments conducted in naïve mice and a Xenogen IVIS system for experiments conducted in ACLT-induced osteoarthritic joints. For mice receiving HCAd-CMV-Luc, bioluminescence imaging was first performed 24 hours after intra-articular injection, followed by additional imaging at 3 days and 1 week, and then weekly thereafter. D-luciferin (Gold Biotechnology, cat# eLUCK) was prepared at a concentration of 20 mg/mL and administered via intraperitoneal injection at a volume of 100 µL per mouse. Imaging was performed 7 minutes after luciferin injection to allow systemic substrate distribution. Luciferase signal intensity from the knee joint was quantified as total photon flux using Living Image software (Revvity) and used for longitudinal comparison of transgene expression.

### X-gal staining

X-gal staining was performed to assess transgene expression following intra-articular administration of HCAd-CMV-LacZ. Knee joints were harvested 3 days after vector injection and fixed in a fixation solution containing 2% PFA and 0.5% glutaraldehyde (Thomas Scientific, cat# C762Q38) for 20 hours. Fixed tissues were decalcified in 14% EDTA for 3 days, embedded in optimal cutting temperature compound, and cryosectioned at a thickness of 14 µm. Cryosections were incubated in X-gal staining solution (Goldbio, cat# X4281C) at 37 °C for 10 minutes to visualize β-galactosidase activity. Sections were subsequently counterstained with 0.1% Nuclear Fast Red (Thomas Scientific, cat# NC0091942).

### Statistical analysis

All data are presented as mean ± standard deviation (SD), unless otherwise indicated. Statistical analyses were performed using GraphPad Prism software. For comparisons between two groups, unpaired two-tailed Student’s t tests were used. For comparisons involving more than two groups, one-way analysis of variance (ANOVA) was applied as appropriate, followed by post hoc multiple-comparison tests. For analysis of OARSI and synovitis scores, which are ordinal in nature, Kruskal–Wallis test was used, followed by Dunn’s multiple comparisons test. A value of *P* < 0.05 was considered statistically significant.

## Acknowledgements

This work was supported by the U.S. Department of Defense Congressionally Directed Medical Research Programs (CDMRP) (Award No. W81XWH-22-1-0372) and the Lawrence Family Bone Disease Program of Texas. Research reported in this publication was also supported in part by the NICHD’S IDDRC under Award Number P50HD103555 for use of the Rodent Neurobehavior Core. This work was also supported by the RE-JOIN consortium, an NIH HEAL-funded team science initiative to identify the biological underpinnings of chronic joint pain^39^. The RE-JOIN consortium consists of: Armen Akopian, Kyle Allen, Alejandro Almarza, Benjamin Arenkiel, Yangjin Bae, Bruna Balbino de Paula, Anita Bandrowski, Mario Danilo Boada, Jacqueline Boccanfuso, Jyl Boline, Dawen Cai, Dellina Carpio, Robert Caudle, Racel Cela, Yong Chen, Rui Chen, Brian Constantinescu, Cortez, Ibdanelo, Yenisel Cruz-Almeida, M. Franklin Dolwick, Chris Donnelly, Zelong Dou, Joshua Emrick, Malin Ernberg, Danielle Freburg-Hoffmeister, Spencer Fullam, Janak Gaire, Akash Gandhi, Benjamin Goolsby, Stacey Greene, Nele Haelterman, Michael Iadarola, Shingo Ishihara, Azeez Ishola, Sudhish Jayachandran, Zixue Jin, Frank Ko, Priya Kulkarni, Zhao Lai, Brendan Lee, Yona Levites, Carolina Leynes, Jun Li, Martin Lotz, Lindsey Macpherson, Tristan Maerz, Camilla Majano, Anne-Marie Malfait, Maryann Martone, Bella Mehta, Richard Miller, Rachel Miller, Michael Newton, Alia Obeidat, Merissa Olmer, Dana Orange, Miguel Otero, Kevin Otto, Folly Patterson, Marlena Pela, Sienna Perry, Theodore Price, Hernan Prieto, Russell Ray, Dongjun Ren, Margarete Ribeiro Dasilva, Alexus Roberts, Elizabeth Ronan, Oscar Ruiz, Shad Smith, Mairobys Socorro, Kaitlin Southern, Joshua Stover, Michael Strinden, Hannah Swahn, Evelyne Tantry, Sue Tappan, Luis Tovias Sanchez, Airam Vivanco-Estela, Joost Wagenaar, Lai Wang, Kim Worley, Joshua Wythe, and Jiansen Yan.

We acknowledge the Baylor College of Medicine Optical Imaging & Vital Microscopy Core (OIVM) for assistance with phase-contrast µCT imaging. All schematic illustrations were generated using BioRender.com.

## Author contributions

J.Y., M.S., Y.B. and B.H.L.L. designed the study; J.Y., Z.D., C.F.M., R.C., M.M.J., S.S.M., J.A.A, D.R.C, A.R.S., L.A.Y., O.E.R, D.J.P., P.N. performed the experiments; J.Y., Z.D., S.V., O.E.R, P.N., N.A.H, M.S., Y.B., B.H.L.L. analyzed and discussed the results; J.Y. wrote the draft; N.A.H., M.S., Y.B., B.H.L.L. edited the manuscript. All authors reviewed and approved the final manuscript.

## Declaration of generative AI and AI-assisted technologies in the manuscript preparation process

The authors wrote the original draft of this manuscript. During the preparation of this work, the authors used OpenAI to improve language and readability. After using this tool, the authors reviewed and edited the content as needed and take full responsibility for the content of the published article.

## References

1. Kloppenburg, M., Namane, M., and Cicuttini, F. (2025). Osteoarthritis. Lancet 405, 71–85. 10.1016/S0140-6736(24)02322-5.

2. Jenei-Lanzl, Z., Meurer, A., and Zaucke, F. (2019). Interleukin-1β signaling in osteoarthritis - chondrocytes in focus. Cell Signal 53, 212–223. 10.1016/j.cellsig.2018.10.005.

3. Mailhot, B., Christin, M., Tessandier, N., Sotoudeh, C., Bretheau, F., Turmel, R., Pellerin, È., Wang, F., Bories, C., Joly-Beauparlant, C., et al. (2020). Neuronal interleukin-1 receptors mediate pain in chronic inflammatory diseases. J Exp Med 217, e20191430. 10.1084/jem.20191430.

4. Chevalier, X., Goupille, P., Beaulieu, A.D., Burch, F.X., Bensen, W.G., Conrozier, T., Loeuille, D., Kivitz, A.J., Silver, D., and Appleton, B.E. (2009). Intraarticular injection of anakinra in osteoarthritis of the knee: a multicenter, randomized, double-blind, placebo-controlled study. Arthritis Rheum 61, 344–352. 10.1002/art.24096.

5. Aitken, D., Cai, G., Hill, C.L., Cicuttini, F.M., Wluka, A.E., Wang, Y., Keen, H.I., Thompson, M.J.W., Asthana, C., Antony, B.E., et al. (2026). Diacerein for Knee Osteoarthritis: A Randomized Clinical Trial. JAMA Intern Med, e258237. 10.1001/jamainternmed.2025.8237.

6. Kloppenburg, M., Peterfy, C., Haugen, I.K., Kroon, F., Chen, S., Wang, L., Liu, W., Levy, G., Fleischmann, R.M., Berenbaum, F., et al. (2019). Phase IIa, placebo-controlled, randomised study of lutikizumab, an anti-interleukin-1α and anti-interleukin-1β dual variable domain immunoglobulin, in patients with erosive hand osteoarthritis. Ann Rheum Dis 78, 413–420. 10.1136/annrheumdis-2018-213336.

7. Grol, M.W., and Lee, B.H. (2018). Gene therapy for repair and regeneration of bone and cartilage. Curr Opin Pharmacol 40, 59–66. 10.1016/j.coph.2018.03.005.

8. Evans, C.H., Ghivizzani, S.C., Keravala, A., Chalberg, T.W., and Robbins, P.D. (2025). Osteoarthritis gene therapy: Expanding the scope of genetic therapies. Mol Ther 33, 3456–3457. 10.1016/j.ymthe.2025.07.019.

9. Rosewell, A., Vetrini, F., and Ng, P. (2011). Helper-Dependent Adenoviral Vectors. J Genet Syndr Gene Ther Suppl 5, 001. 10.4172/2157-7412.s5-001.

10. Hackett, N.R., and Crystal, R.G. (2025). Four decades of adenovirus gene transfer vectors: History and current use. Mol Ther 33, 2192–2204. 10.1016/j.ymthe.2025.03.062.

11. Stone, A., Grol, M.W., Ruan, M.Z.C., Dawson, B., Chen, Y., Jiang, M.-M., Song, I.- W., Jayaram, P., Cela, R., Gannon, F., et al. (2019). Combinatorial Prg4 and Il-1ra Gene Therapy Protects Against Hyperalgesia and Cartilage Degeneration in Post-Traumatic Osteoarthritis. Hum Gene Ther 30, 225–235. 10.1089/hum.2018.106.

12. Senter, R., Boyce, R., Repic, M., Martin, E.W., Chabicovsky, M., Langevin-Carpentier, G., Bédard, A., and Bodick, N. (2022). Efficacy and Safety of FX201, a Novel Intra-Articular IL-1Ra Gene Therapy for Osteoarthritis Treatment, in a Rat Model. Hum Gene Ther 33, 541–549. 10.1089/hum.2021.131.

13. Nixon, A.J., Grol, M.W., Lang, H.M., Ruan, M.Z.C., Stone, A., Begum, L., Chen, Y., Dawson, B., Gannon, F., Plutizki, S., et al. (2018). Disease-Modifying Osteoarthritis Treatment With Interleukin-1 Receptor Antagonist Gene Therapy in Small and Large Animal Models. Arthritis Rheumatol 70, 1757–1768. 10.1002/art.40668.

14. Cohen, S., Conaghan, P., Hochberg, M., Kivitz, A., Kim, M., Joy, N., Jiang, S., DiGiorgi, M., Slonin, J., and Slonin, D. (2025). PCRX-201 high-capacity adenovirus serotype 5 gene therapy demonstrates sustained clinical efficacy and safety in patients with knee osteoarthritis [abstract]. Arthritis Rheumatol. 77 (suppl 9).

15. Bulaklak, K., and Gersbach, C.A. (2020). The once and future gene therapy. Nat Commun 11, 5820. 10.1038/s41467-020-19505-2.

16. Ruan, M.Z.C., Erez, A., Guse, K., Dawson, B., Bertin, T., Chen, Y., Jiang, M.-M., Yustein, J., Gannon, F., and Lee, B.H.L. (2013). Proteoglycan 4 expression protects against the development of osteoarthritis. Sci Transl Med 5, 176ra34. 10.1126/scitranslmed.3005409.

17. Wang, W.-C., Sayedahmed, E.E., and Mittal, S.K. (2022). Significance of Preexisting Vector Immunity and Activation of Innate Responses for Adenoviral Vector-Based Therapy. Viruses 14, 2727. 10.3390/v14122727.

18. Sumida, S.M., Truitt, D.M., Lemckert, A.A.C., Vogels, R., Custers, J.H.H.V., Addo, M.M., Lockman, S., Peter, T., Peyerl, F.W., Kishko, M.G., et al. (2005). Neutralizing antibodies to adenovirus serotype 5 vaccine vectors are directed primarily against the adenovirus hexon protein. J Immunol 174, 7179–7185. 10.4049/jimmunol.174.11.7179.

19. Crawford-Miksza, L., and Schnurr, D.P. (1996). Analysis of 15 adenovirus hexon proteins reveals the location and structure of seven hypervariable regions containing serotype-specific residues. J Virol 70, 1836–1844. 10.1128/jvi.70.3.1836-1844.1996.

20. Mack, C.A., Song, W.R., Carpenter, H., Wickham, T.J., Kovesdi, I., Harvey, B.G., Magovern, C.J., Isom, O.W., Rosengart, T., Falck-Pedersen, E., et al. (1997). Circumvention of anti-adenovirus neutralizing immunity by administration of an adenoviral vector of an alternate serotype. Hum Gene Ther 8, 99–109. 10.1089/hum.1997.8.1-99.

21. Goodrich, L.R., Grieger, J.C., Phillips, J.N., Khan, N., Gray, S.J., McIlwraith, C.W., and Samulski, R.J. (2015). scAAVIL-1ra dosing trial in a large animal model and validation of long-term expression with repeat administration for osteoarthritis therapy. Gene Ther 22, 536–545. 10.1038/gt.2015.21.

22. Loeb, E.J., Fergione, S.A., Yudistyra, V., Fanous, M.M., Benkert, A.R., Fisher, D.G., Hull, J.A., ElMallah, M.K., and Asokan, A. (2025). Complete neutralizing antibody evasion by serodivergent non-mammalian AAVs enables gene therapy redosing. Cell Rep Med 6, 102475. 10.1016/j.xcrm.2025.102475.

23. Roberts, D.M., Nanda, A., Havenga, M.J.E., Abbink, P., Lynch, D.M., Ewald, B.A., Liu, J., Thorner, A.R., Swanson, P.E., Gorgone, D.A., et al. (2006). Hexon-chimaeric adenovirus serotype 5 vectors circumvent pre-existing anti-vector immunity. Nature 441, 239–243. 10.1038/nature04721.

24. Doshi, B.S., Markmann, C.A., Novak, N., Juarez Rojas, S., Davidson, R., Chau, J.Q., Wang, W., Carrig, S., Martos Rus, C., Samelson-Jones, B.J., et al. (2025). Use of CD19-targeted immune modulation to eradicate AAV-neutralizing antibodies. Mol Ther 33, 3073–3085. 10.1016/j.ymthe.2025.03.003.

25. De la Vega, R.E., Sellon, J.L., Smith, J., Wisniewski, S.J., Jurisson, M.L., Frick, M.A., Scrabeck, T.L., Block, J.B., Mills, C.J., Pohlkamp, Z.W., et al. (2025). A phase 1 clinical trial shows safe, sustained, AAV-mediated expression of IL-1Ra in the human osteoarthritic knee joint. Sci Transl Med 17, eadu9804. 10.1126/scitranslmed.adu9804.

26. Gouze, J.-N., Gouze, E., Palmer, G.D., Liew, V.S., Pascher, A., Betz, O.B., Thornhill, T.S., Evans, C.H., Grodzinsky, A.J., and Ghivizzani, S.C. (2003). A comparative study of the inhibitory effects of interleukin-1 receptor antagonist following administration as a recombinant protein or by gene transfer. Arthritis Res Ther 5, R301–309. 10.1186/ar795.

27. Greene, M.A., and Loeser, R.F. (2015). Aging-related inflammation in osteoarthritis. Osteoarthritis Cartilage 23, 1966–1971. 10.1016/j.joca.2015.01.008.

28. Yang, Y.-S., Kim, M.-J., Chaugule, S., Mayer, E., DeSouza, N., Ma, H., Xie, J., Lee, K.-Y., Li, S., Gravallese, E., et al. (2025). Nature-inspired IL-1 targeted therapy to treat chronic inflammatory diseases. Mol Ther 33, 6379–6397. 10.1016/j.ymthe.2025.09.008.

29. Malfait, A.-M., and Schnitzer, T.J. (2013). Towards a mechanism-based approach to pain management in osteoarthritis. Nat Rev Rheumatol 9, 654–664. 10.1038/nrrheum.2013.138.

30. Lee, N.K., and Chang, J.W. (2024). Manufacturing Cell and Gene Therapies: Challenges in Clinical Translation. Ann Lab Med 44, 314–323. 10.3343/alm.2023.0382.

31. Zhou, K., Yuan, M., Sun, J., Zhang, F., Li, X., Xiao, X., and Wu, X. (2025). Co-delivery of IL-1Ra and SOX9 via AAV inhibits inflammation and promotes cartilage repair in surgically induced osteoarthritis animal models. Gene Ther 32, 211–222. 10.1038/s41434-025-00515-y.

32. Chen, B., Qin, J., Wang, H., Magdalou, J., and Chen, L. (2010). Effects of adenovirus-mediated bFGF, IL-1Ra and IGF-1 gene transfer on human osteoarthritic chondrocytes and osteoarthritis in rabbits. Exp Mol Med 42, 684–695. 10.3858/emm.2010.42.10.067.

33. Suzuki, M., Cela, R., Clarke, C., Bertin, T.K., Mouriño, S., and Lee, B. (2010). Large-scale production of high-quality helper-dependent adenoviral vectors using adherent cells in cell factories. Hum Gene Ther 21, 120–126. 10.1089/hum.2009.096.

34. Glasson, S.S., Blanchet, T.J., and Morris, E.A. (2007). The surgical destabilization of the medial meniscus (DMM) model of osteoarthritis in the 129/SvEv mouse. Osteoarthritis Cartilage 15, 1061–1069. 10.1016/j.joca.2007.03.006.

35. Morita, D., Rosewell Shaw, A., Biegert, G., Porter, C., Woods, M., Vasileiou, S., Lim, B., and Suzuki, M. (2024). Additional expression of T-cell engager in clinically tested oncolytic adeno-immunotherapy redirects tumor-infiltrated, irrelevant T cells against cancer cells to enhance antitumor immunity. J Immunother Cancer 12, e009741. 10.1136/jitc-2024-009741.

36. Glasson, S.S., Chambers, M.G., Van Den Berg, W.B., and Little, C.B. (2010). The OARSI histopathology initiative - recommendations for histological assessments of osteoarthritis in the mouse. Osteoarthritis Cartilage 18 *Suppl 3*, S17–23. 10.1016/j.joca.2010.05.025.

37. Obeidat, A.M., Kim, S.Y., Burt, K.G., Hu, B., Li, J., Ishihara, S., Xiao, R., Miller, R.E., Little, C., Malfait, A.-M., et al. (2024). A standardized approach to evaluation and reporting of synovial histopathology in two surgically induced murine models of osteoarthritis. Osteoarthritis Cartilage 32, 1273–1282. 10.1016/j.joca.2024.05.006.

38. Ruan, M.Z.C., Dawson, B., Jiang, M.-M., Gannon, F., Heggeness, M., and Lee, B.H.L. (2013). Quantitative imaging of murine osteoarthritic cartilage by phase-contrast micro-computed tomography. Arthritis Rheum 65, 388–396. 10.1002/art.37766.

39. Haelterman, N.A., Akopian, A.N., Allen, K.D., Cruz-Almeida, Y., Donnelly, C.R., Lee, B., Lenzi, R., Malfait, A.-M., Mancini, M., Martone, M.E., et al. Designing and implementing solution-oriented team science initiatives—a chronic pain example. Front Pain Res (Lausanne) 6, 1669072. 10.3389/fpain.2025.1669072.

