## Supplemental figures for "Sequential intra-articular HCAd-NFκB-IL-1Ra delivery improves therapeutic efficacy in post-traumatic osteoarthritis"

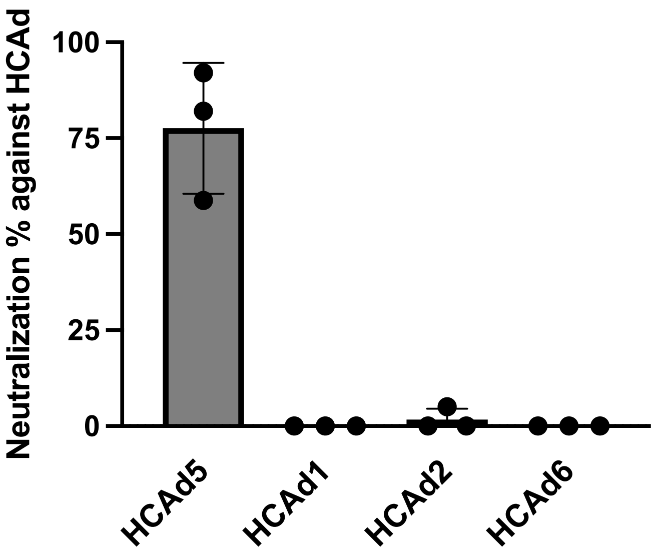


**Figure S1. Neutralizing antibody responses against HCAd serotypes following FGAd5 immunization.**
Serum was collected one week after two intramuscular immunizations with FGAd5 and incubated with different HCAd-CMV-Luc reporter vectors (HCAd5, HCAd1, HCAd2, and HCAd6) prior to infection of A549 cells. Neutralization was assessed based on reporter gene expression in target cells and normalized to no-serum controls. Serum from FGAd5-immunized efficiently neutralized HCAd5 but showed no detectable neutralizing activity against HCAd1, HCAd2, or HCAd6 (n = 3).

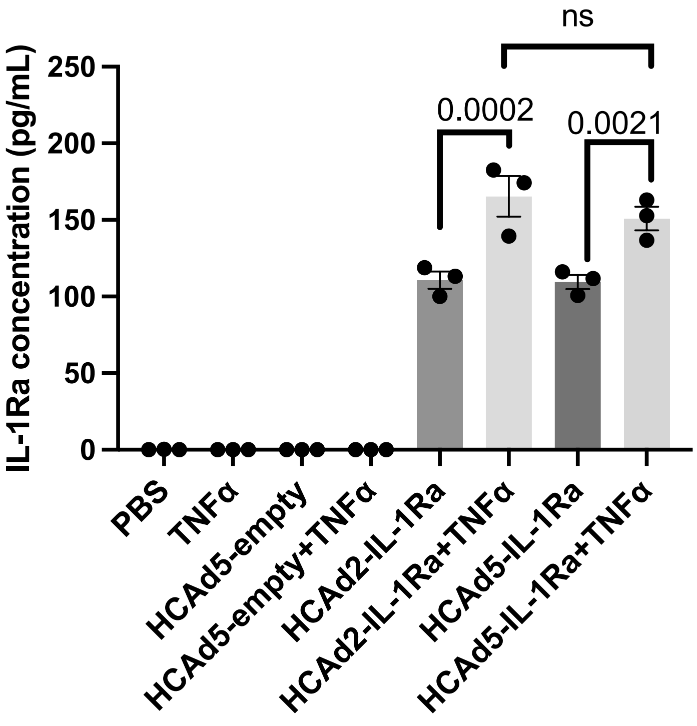


**Figure S2. Comparable IL-1Ra expression by HCAd5 and HCAd2 under basal and inflammatory conditions in vitro.**
HEK293 cells were transduced with HCAd5-NFκB-IL-1Ra or HCAd2-NFκB-IL-1Ra and subsequently stimulated with TNFα. IL-1Ra levels in the culture supernatant were quantified by ELISA. Both vectors produced comparable IL-1Ra at baseline and showed similar induction following TNFα stimulation (n = 3).
